# Identification of HLA-A33-restricted CD8^+^ T cell epitopes from avian influenza A/H5N1

**DOI:** 10.64898/2026.06.21.733083

**Authors:** A.K.M. Muraduzzaman, Patricia T. Illing, Mitchell Jenzen, Nathan P. Croft, Sinead M. Williams, Paul Selleck, Michelle L. Baker, Katherine Kedzierska, Anthony W. Purcell, Nicole A. Mifsud

## Abstract

The rapid evolution of avian influenza A/H5N1, including the recent U.S. clade 2.3.4.4b outbreak, highlights its pandemic potential and the urgent need for durable, broadly protective vaccines. Given the capacity of CD8^+^ T cells to mediate cross-strain immunity, we investigated whether geographically distinct HLA-A33 allotypes, HLA-A*33:01 in East/Southeast Asia and HLA-A*33:03 in South Asia, differentially shape the influenza immunopeptidome and influence antiviral immunity. Antigen-presenting cells overexpressing HLA-A*33:01 or HLA-A*33:03 were transfected with single A/H5N1 antigens or infected with A/X-31 (H3N2) as a control comparison representing current seasonal influenza virus. We identified novel ligands restricted to HLA-A*33:01 (57 from A/H5N1; 55 from A/X-31) and HLA-A*33:03 (29 from A/H5N1; 45 from A/X-31). Although fewer peptides were recovered for HLA-A*33:03, a larger proportion of A/X-31-derived peptides were predicted as high-affinity binders (74%) compared with HLA-A*33:01 (61%), indicating qualitative differences in antigen presentation. To determine immunogenicity, peripheral blood lymphocytes from HLA-A*33:03-positive, A/H5N1-naïve donors were stimulated with four conserved peptides: PB2_GTF_, PB2_KTY_, NP_SVQ_ and PB1_MTK_. All elicited robust CD8⁺ T cell activation despite the absence of prior A/H5N1 exposure, demonstrating cross-recognition by memory T cells primed against seasonal influenza. These findings define HLA-A33-restricted influenza epitopes and reveal allotype-specific presentation features that shape CD8^+^ T cell immunity. Conserved, immunogenic peptides identified here represent promising candidates for rational design of broadly cross-reactive vaccines to protect HLA-A33-expressing populations against severe A/H5N1 disease. Data are available via ProteomeXchange with identifier PXD078870.

**Author Summary:** Avian influenza A/H5N1 continues to pose a significant pandemic threat because of its ability to infect humans and its potential to acquire sustained human-to-human transmissibility. While current influenza vaccines primarily target rapidly evolving viral surface proteins, CD8^+^ T cells can recognize more conserved internal viral proteins and may provide broader protection against diverse influenza strains.

In this study, we investigated how two common HLA-A33 variants, which are prevalent in South, East, and Southeast Asian populations, present influenza-derived peptides to CD8^+^ T cells. We identified novel influenza peptides presented by HLA-A*33:01 and HLA-A*33:03. Importantly, several conserved A/H5N1-derived peptides were recognized by memory CD8^+^ T cells from healthy individuals with no prior exposure to A/H5N1, suggesting that previous infection with seasonal influenza viruses can generate cross-reactive immune responses.

Our findings expand the current repository of influenza T cell targets and provide new insights into antiviral immunity in HLA-A33-expressing populations. The conserved and immunogenic peptides identified in this study may help guide the development of broadly protective influenza vaccines and contribute to future pandemic preparedness efforts.

## Introduction

Influenza A virus (IAV) is a ubiquitous respiratory pathogen responsible for annual seasonal epidemics and periodic pandemics in humans, with substantial global mortality and economic burden [1]. Contemporary seasonal strains comprise two IAV subtypes (A/H1N1pdm09 and A/H3N2) and two lineages of influenza B viruses (B/Victoria and B/Yamagata) [2]. Although the B/Yamagata lineage has not circulated since 2020 and is now considered extinct [3]. Beyond humans, IAVs infect diverse animal hosts, including pigs, horses, marine mammals and birds [4]. IAVs circulating in birds are referred to as avian influenza [5] and carry all IAV subtypes (haemagglutinin [HA1-17] and neuraminidase [NA1-9]) [6, 7]. In recent decades, several new avian IAV strains have emerged at the human-animal interface (*i.e.* H5N6, H5N1, H7N9, H5N8, H9N2, H7N7, H7N4) with different rates of human morbidity and mortality (up to 60-80% for H5N1 and H5N6) [8].

Current vaccines primarily elicit antibodies against the rapidly evolving HA and NA glycoproteins, limiting cross-strain protection [9]. In contrast, internal proteins including nucleoprotein (NP), polymerase acidic (PA), polymerase basic 1 (PB1) and 2 (PB2), matrix 1 (M1), non-structural protein 1 (NS1) and 2 (NS2) are highly conserved, making them attractive targets for long-lasting broader CD8⁺ T cell mediated immunity [10]. Evidence in both humans [11–13] and mice [14–16] demonstrate that cytotoxic T lymphocyte (CTL) killing of influenza infected cells is critical for maintaining immunity and limiting disease pathogenesis through viral clearance [17, 18].

The repertoire of peptides presented by human leukocyte antigen (HLA) class I molecules, known as the immunopeptidome [19], is shaped by extensive HLA polymorphism [20]. This allelic variation largely evolved in response to diverse pathogenic pressure to effectively present peptide antigens on the surface of infected cells [21]. However, not all immune responses are equal, with both qualitative and quantitative differences between HLA variants being reported. For instance, HLA heterozygosity can be advantageous, enabling individuals to present a broader array of viral peptides that results in more effective CD8⁺ T cell responses [22]. Other qualitative features of HLA molecules also shape immune outcomes, including specific peptide-binding motifs, haplotypic context [23], and preferences for peptide ligand length [24], all of which contribute to the establishment of an immunodominance hierarchy [25]. For instance, Quiñones-Parra *et al*., investigated the role of HLA diversity in influenza pathogenesis and reported that HLA-A*03:01, HLA-B*08:01, HLA-B*18:01, HLA-B*27:05 and HLA-B*57:01-specific NP epitopes elicited robust CTL responses, whereas HLA-A*01:01, HLA-A*24:02, HLA-A*68:01 and HLA-B*15:01 showed limited CTL responses in the ethnic groups in which they predominate [14].

In 1996, the highly pathogenic avian influenza A/H5N1 virus (hereafter A/H5N1) emerged in China, with the first human case reported in Hong Kong the following year [26, 27]. Since then, A/H5N1 has spread to 24 countries, resulting in 964 confirmed human cases with a 48% fatality rate [28]. A/H5N1 remains endemic in many Asian regions, particularly in Central, East, South and Southeast Asia [29, 30]. Several factors have been suggested to contribute to the sustained transmission in these countries, including high population density, live bird market practices, climate conditions and host genetics [31–33]. Notably for the latter, HLA allotypes prevalent in different geographical populations influence disease susceptibility and severity [33–35]. As of December 2024, among the Southeast Asian countries, Indonesia reported the highest mortality rate to A/H5N1 infection of 84%, followed by Thailand (68%) and Cambodia (66%), whilst in East Asia, China recorded the highest mortality rate at 58% [28]. Interestingly, despite widespread environmental contamination, particularly in live bird markets [30, 36], the incidence and mortality rates of human infection in South Asian populations remain significantly lower (2.5-fold), with an average case fatality rate of 28% compared to 69% reported in other regions of Asia [28].

Given this lower susceptibility and mortality of A/H5N1 in South Asia, this study explored CD8^+^ T cell epitopes restricted to the most prevalent HLA class I (HLA-I) alleles in the region. According to the Allele Frequency Net Database [37], HLA-A*11:01 (gene frequency [gf] of 13%), HLA-A*24:02 (gf 12.5%) and HLA-A*33:03 (gf 10.5%) are the most common HLA-A allotypes in Asian populations. Having already characterised CD8^+^ T cell epitopes from human IAV and IBV in HLA-A*24:02 and HLA-A*11:01 allotypes [38, 39], we focused on HLA-A*33:03. This allotype is predominantly expressed in South Asian populations and we compared antigen presentation with the closely related HLA-A*33:01 allele, more commonly found in East and Southeast Asian countries (gf 11.7% in Han Chinese). HLA-A*33:01 and HLA-A*33:03 differ by two amino acid residues at positions 171 (H171Y) and 186 (R186K), which are likely to influence both seasonal influenza A/X-31 and avian influenza A/H5N1 peptide repertoires and CD8^+^ T cell recognition that contribute to the protective HLA-A*33:03-specific immunity observed in South Asian populations.

## Results

### Differential peptide presentation by HLA-A33 allotypes following A/X-31 infection

To define how closely related HLA-A33 allotypes shape influenza antigen presentation, we performed immunopeptidomic profiling of C1R parental, C1R.A*33:01, and C1R.A*33:03 cells following a 12 hr A/X-31 (H3N2) infection. Infection efficiency was observed to be 32% in C1R.A*33:01 cells and 23% in C1R.A*33:03 cells (**Figure 1A**). Across both mock and infected conditions (biological duplicates), 12 datasets were generated. Peptides identified in parental cells (co-express normal levels of HLA-B*35:03 and HLA-C*04:01) were excluded from HLA-A33 analyses.

**Figure 1.**
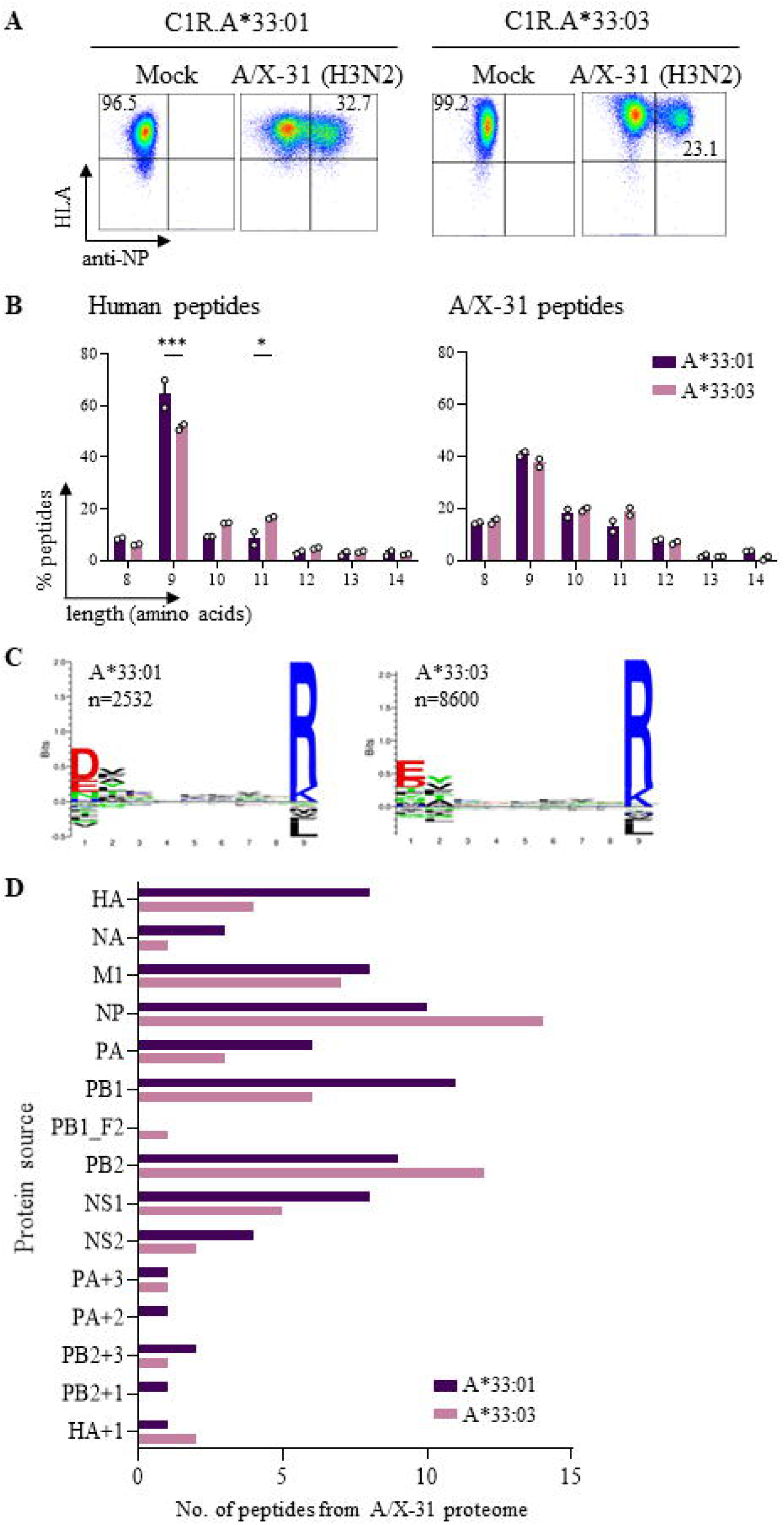
LC-MS/MS analysis of HLA-A*33 restricted A/X-31 influenza peptides. **A.** Dot plot showing A/X-31 infection efficiency in C1R transfected cells after 12 hrs. **B.** Ligands isolated from C1R.A*33:03 and C1R.A*33:01 using immunoaffinity capture with pan HLA-I antibody W6/32. Human peptides (8-12 amino acids) and influenza peptides (8-14 amino acids) were analysed at 1% FDR after removing the C1R parental background. Each experiment contains two biological replicates per condition. Two-way ANOVA followed by Sidak’s MCT was performed for statistical analysis of data [*p<0.05, ***p<0.001] (GraphPad Prism 9.0.0, USA). **C.** Binding motif analysis of human 9mer peptides using Seq2logo2.0 [82] (Note: Motifs were identical in both infected and uninfected datasets; therefore, only those from the uninfected dataset are presented here.) **D.** Distribution of A/X-31-derived A*33:03 and A*33:01 ligands across the viral proteomes, including alternate reading frame (donated as + after protein name).

Over 95% of peptides were 9-11 amino acids in length, dominated by 9mers that accounted for 60% of human and 40% of A/X-31-derived peptides. Notably, HLA-A*33:03 presented fewer human 9mers and more 11mers compared to HLA-A*33:01, with a similar but non-significant trend observed for A/X-31-derived peptides (**Figure 1B**), indicating subtle but consistent differences in peptide length preferences. Both allotypes displayed highly similar 9mer peptide-binding motifs, with enrichment of phenylalanine (F), valine (V), threonine (T) or alanine (A) at position 2 (P2), and arginine (R) or lysine (K) at the C-terminal anchor (P9). Differences were observed at position 1 (P1), where aspartic acid (D) was enriched in peptides presented by HLA-A*33:01, whereas glutamic acid (E) was more frequent in HLA-A*33:03. Tyrosine (Y) at P2 was also more enriched in HLA-A*33:03. Apart from these position-specific differences, amino acid usage across other positions was largely comparable between the two allotypes (**Figure 1C; Supplementary Figure 1**). Similar trends were observed for longer peptides (10–12mers) (**Supplementary Figure 2**).

We identified 73 and 58 curated A/X-31-derived peptides presented by HLA-A*33:01 and HLA-A*33:03, respectively (**Supplementary Data 1**). These peptides span the viral proteome (except PB1_F2 for HLA-A*33:01 and M2 for both), including those derived from alternative reading frames that do not map to canonical proteins, denoted by “+” following the canonical protein name (**Figure 1D**).

Despite similar binding motifs, only 22.9% of human-derived peptides (8-12mer) and 31% of A/X-31-derived peptides (8-14mer) were commonly presented by both HLA-A33 allotypes (**Figure 2A, B**). Notably, 70% of human peptides presented by HLA-A*33:03 were classified as binders, whereas 48% of peptides bound to HLA-A*33:01 (**Figure 2C**). Similarly, for A/X-31 peptides, 74% strongly bound to HLA-A*33:03 compared to 61% for HLA-A*33:01 (**Figure 2D**). Importantly, the majority of viral peptides identified were novel, with only 18/73 (24.6%) HLA-A*33:01-restricted peptides and 13/58 (22.4%) HLA-A*33:03-restricted peptides reported in the Immune Epitope Database (IEDB) (**Supplementary Data 1**) [40].

**Figure 2.**
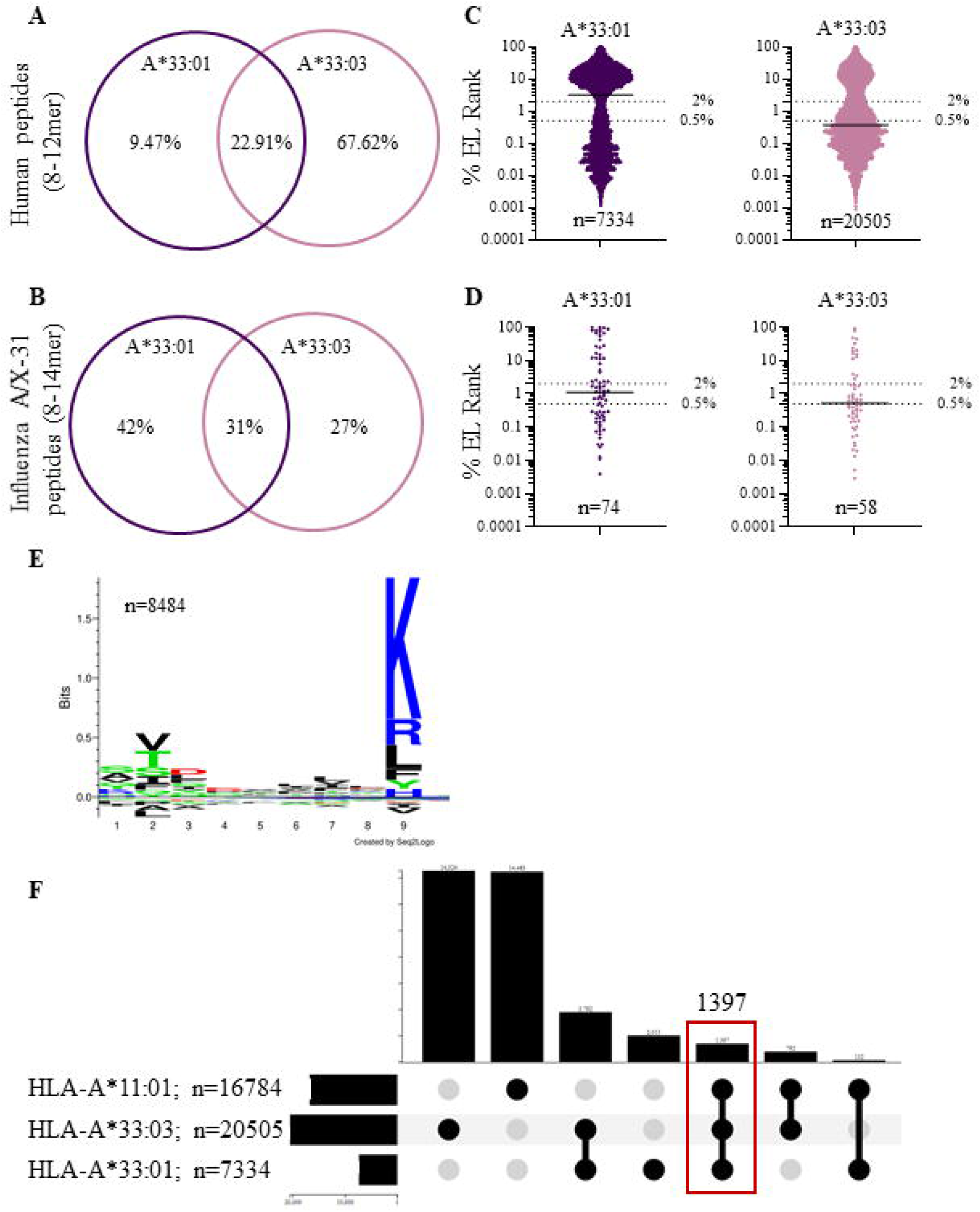
Human and influenza A/X-31 peptides overlapping and predicted binding affinity between A*33:01 and A*33:03. (A, **B**). 70% peptides were unique between A*33:01 and A*33:03. For human peptides only 8-12mers and for A/X-31, 8-14 mers peptides were considered. **C**. Here, 70% of human derived peptides were predicted binders to HLA-A*33:03 compared to 48% binders to HLA-A*33:01. **D**. 74% of A*33:03 restricted A/X-31 peptides were predicted binders compared to 62% of the identified influenza peptides were binders to A*33:01. Peptide-binding affinities were predicted using NetMHCpan-4.2 [44], with % rank determining strong binders (<0.5 SB), weak binders (<2 WB), non-binders (>2 NB). **E**. HLA-A*11:01-restricted human derived 9mer peptides motifs generated by Seq2logo2.0 [82]. **F**. Upset plot showing human peptide overlap among three members of the HLA-A3 supertype family.

HLA supertypes are comprised of different allotypes that share similar peptide binding motifs, with both HLA-A11 and HLA-A33 belonging to the HLA-A3 supertype. Comparative influenza A/X-31 peptide repertoire analysis between this study and our previous report on HLA-A*11:01 [38] revealed marked motif divergence at P1 and P9 (**Figure 2E**) and minimal peptide overlap with only 0.68% viral peptides shared across all three allotypes (**Figure 2F, Supplementary Data 1**), underscoring functional non-redundancy within the HLA-A3 supertype.

### HLA-A33 immunopeptidomes reveal novel A/H5N1-derived peptides

As productive avian influenza A/H5N1 of C1R parental or C1R.A33 expressing cells was not achievable, individual viral proteins (NP, PB2, M1, HA) were expressed in these cells. All A/H5N1 transfected cell lines were bulk sorted for co-expression of HLA-I (GFP+) and influenza (DsRed+) proteins by flow cytometry, achieving >88% purity (**Figure 3A**). For each line, 1 × 10^9^ cells were used for immunoaffinity purification of pHLA-I complexes coupled with LC-MS/MS to identify A/H5N1 protein-derived peptides (8-14mers). As expected, 9-11mers dominated all HLA-I ligands (80% of total peptides), with more than 40% represented by 9mers for both HLA-A33 allotypes (**Figure 3B**). Similar to human peptide data (**Figure 1B**), HLA-A*33:03 mostly presented less 8-9mer peptides and more 10-14mer pepitides in all four A/H5N1 proteins (**Figure 3B**) with both allotypes showing similar peptide binding motifs between natural A/X-31 infection (**Figure 1C**) and A/H5N1 protein-transfected cell lines (**Supplementary Figure 2A**).

**Figure 3.**
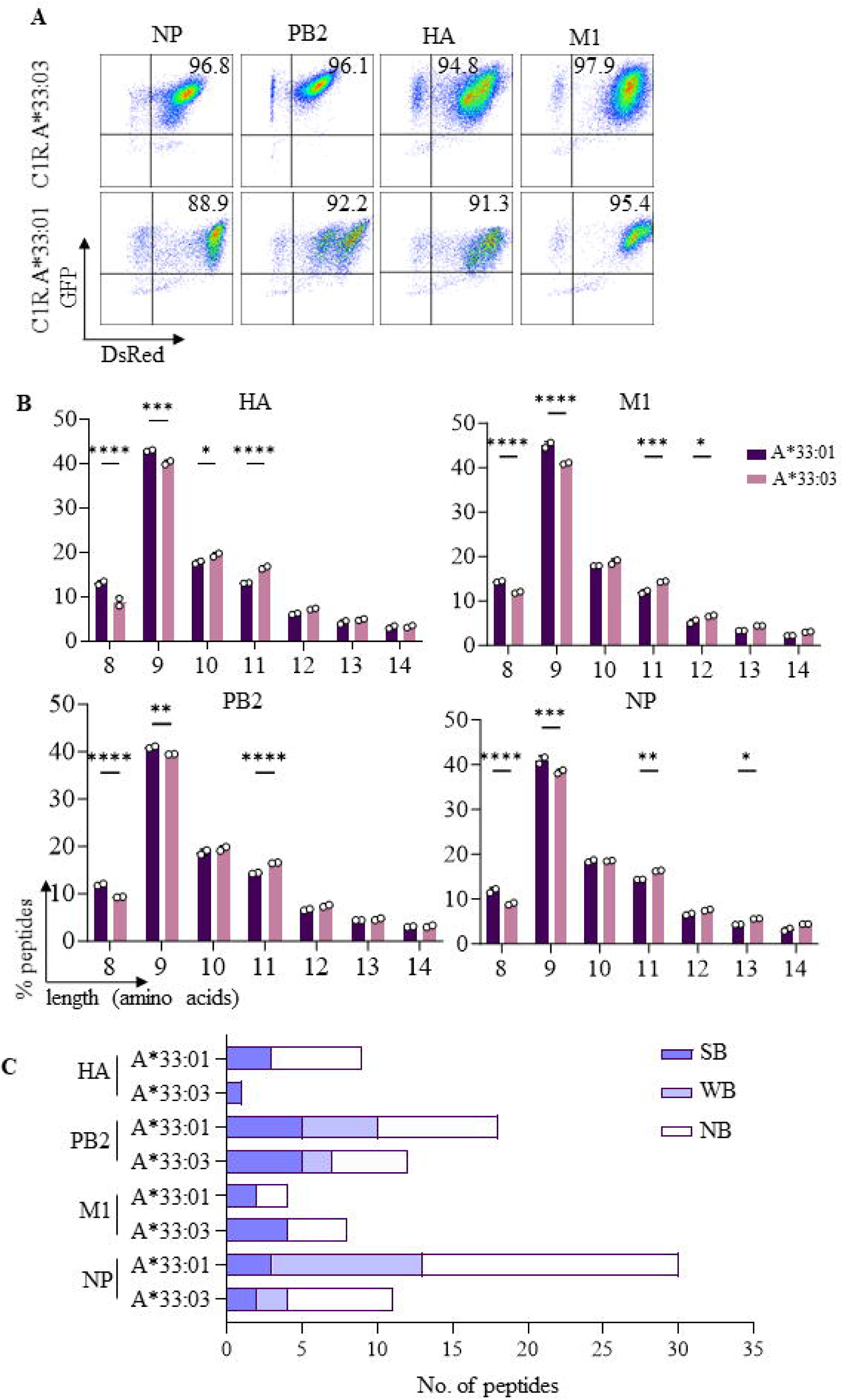
A/H5N1-transfected immunopeptidomics data were comparable to naturally infected A/X-31. **A.** Flow cytometry plots showing individual A/H5N1 protein (HA, M1, PB2, NP) transfection efficiencies, including their HLA expression in C1R transfected cells. **B.** Percentage length distribution of combined human and influenza protein-derived HLA-I ligands of C1R.A*33:03 and C1R.A*33:01 isolated using the pan HLA-I antibody W6/32. 8-14 amino acid long peptides were analysed at 1% FDR after removal of the C1R parental background. Each experiment included two biological replicates. Statistical analysis was performed using Two-way ANOVA followed by Sidak’s MCT [*p<0.05, **p<0.01, ***p<0.001, ****p<0.0001] (GraphPad Prism 9.0.0, USA). **C.** Distribution of A/H5N1 derived peptides from A*33:03 and A*33:01 ligands in C1R cells transfected with individual A/H5N1 viral proteins. Peptide-binding affinities were predicted using NetMHCpan-4.2 [44], with % rank determining strong binders (<0.5 SB), weak binders (<2 WB), non-binders (>2 NB).

A total of 61 (HA=9, NP=30, PB2=18, M1=4) and 32 (HA=1, NP=11, PB2=12, M1=8) A/H5N1-derived peptides were identified for HLA-A*33:01 and HLA-A*33:03, respectively (**Figure 3C**). Srikingly, 57/61 (93%) HLA-A*33:01-restricted and 29/32 (91%) HLA-A*33:03-restricted peptides are unreported in IEDB [40], substantially expanding the known HLA-A33-restricted A/H5N1 epitope repertoire (**Supplementary Data 2**). Predicted binding affinities were comparable between allotypes, with 46% and 50% of peptides classified as binders for HLA-A*33:01 and HLA-A*33:03, respectively, supporting quantitatively similar peptide presentation despite differences in repertoire breadth (**Supplemetary Figure 3 A, B**).

### Identification of immunogenic HLA-A*33:03-restricted CD8^+^ T cell epitopes

Whilst influenza peptides identified by immunopeptidomics are naturally processed and presented, only a subset are recognised by T cells [38, 39, 41–43]. To determine which ligands were immunogenic, peptides from A/X-31 and A/H5N1 datasets were selected based on (i) HLA-restriction, (ii) source protein and (iii) predicted binding affinity [44] (**Table 1**). The top ranked 29 candidate peptides were selected, comprising of 3 HLA-A*33:01-restricted, 9 HLA-A*33:03-restricted, and 17 presented by both allotypes. Among these, 11 peptides were identified in A/H5N1, 7 in A/X-31 and 11 were shared by both A/X-31 and A/H5N1. Three peptides presented by both HLA-A*33 allotypes following infection with A/X-31 and transfection with A/H5N1 antigens, despite lacking predicted HLA binding affinity (*i.e.* non-binder) were also included. Only two peptides have been reported to elicit a positive response in T cell IFNγ assays in IEDB [40]. These 29 peptides were divided initially into six pools for T cell screening, followed by evaluation of single peptides within pools that generated an immune response (**Table 1**).

**Table 1.**
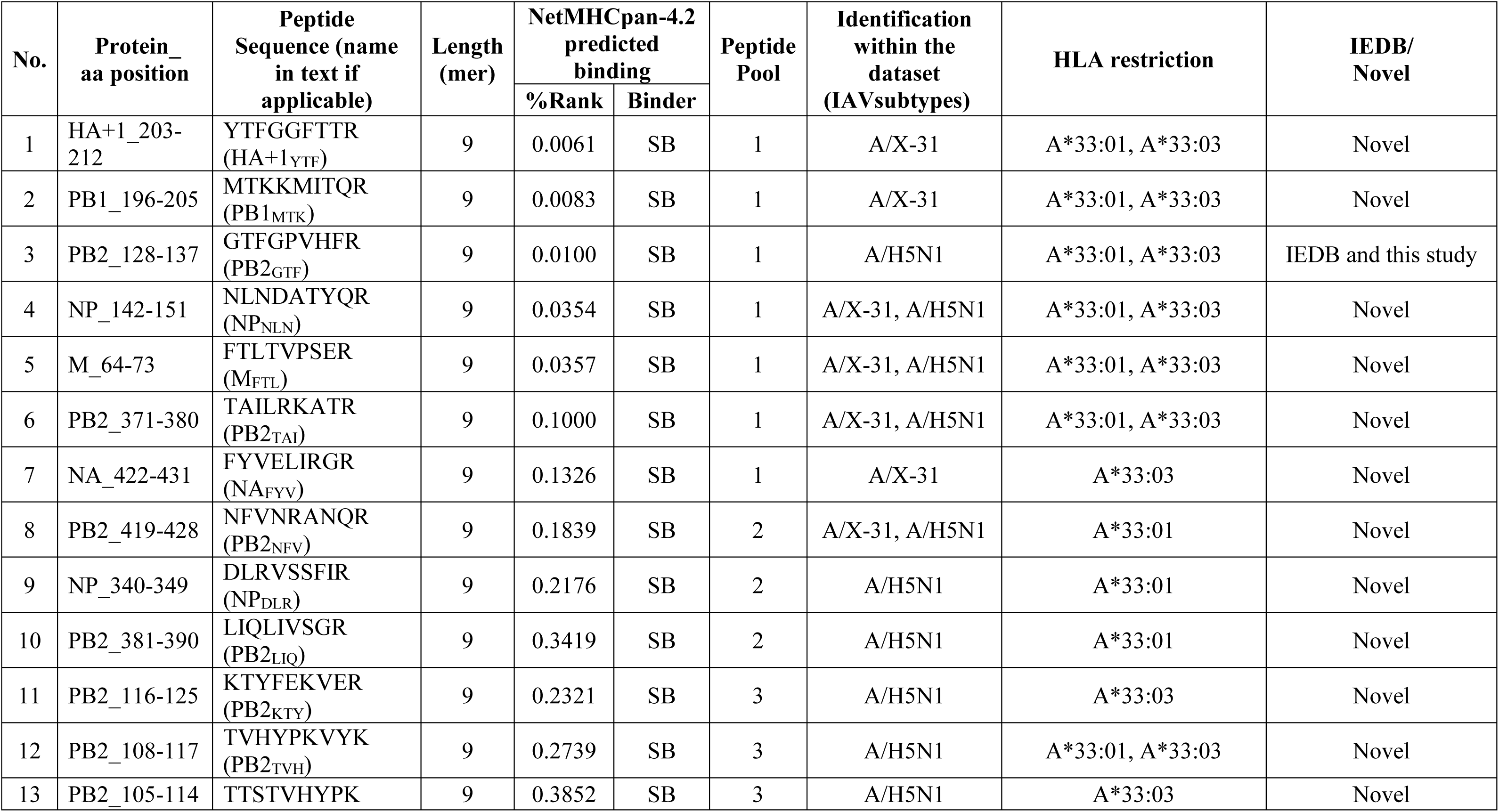
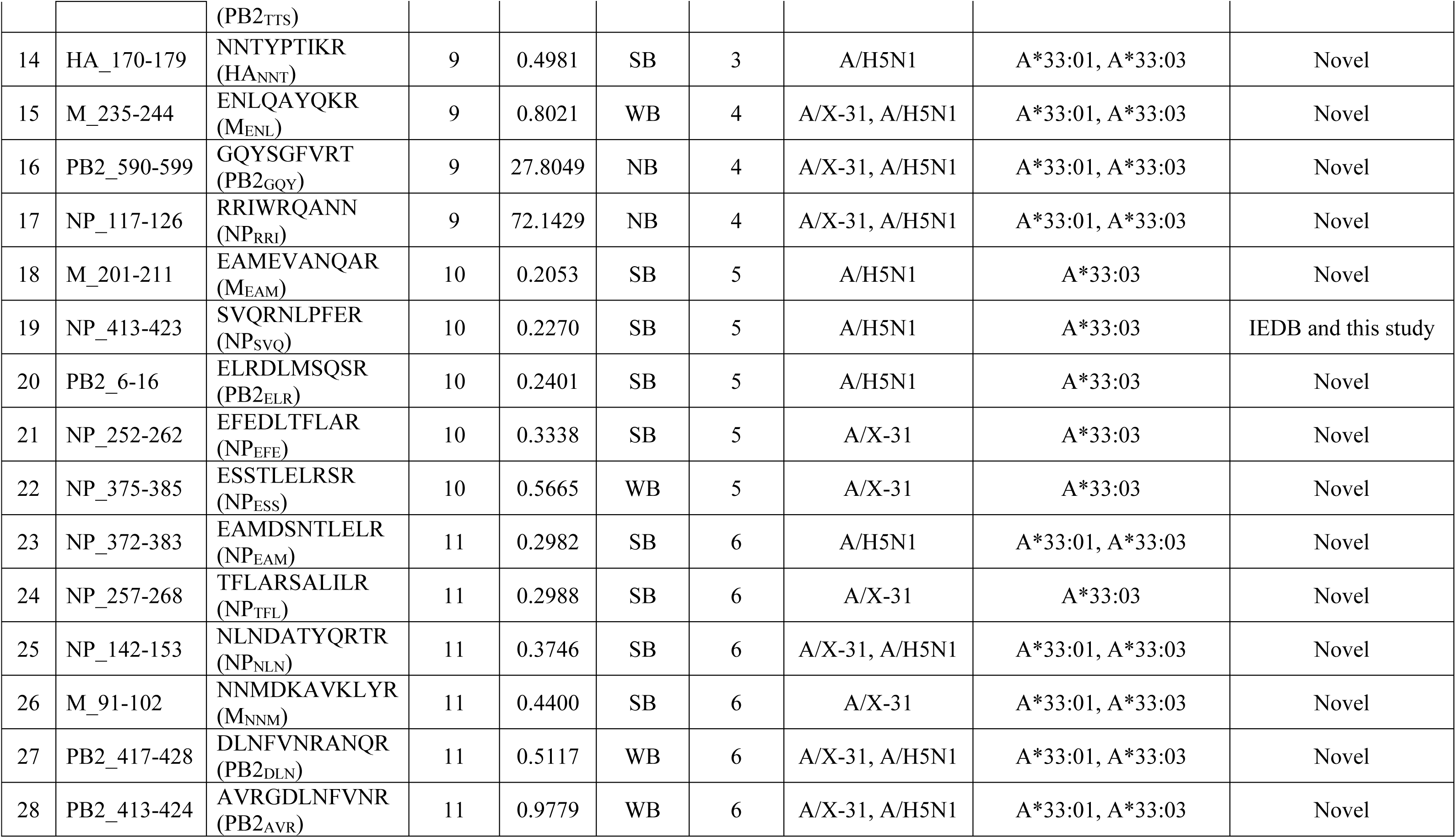
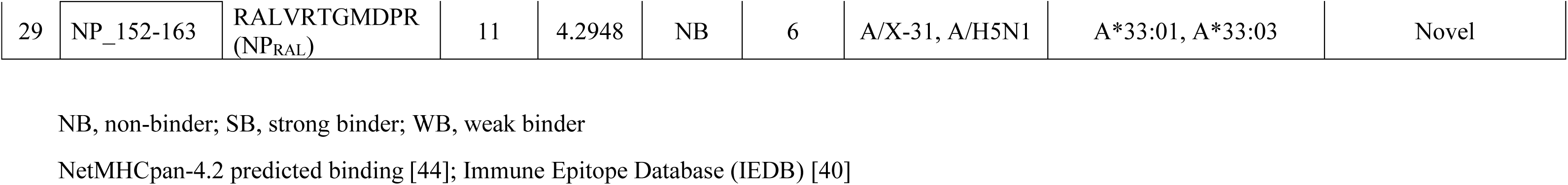
Selected list of naturally presented A/X-31 and A/H5N1 peptides evaluated for T cell recognition.

Briefly, influenza-specific T cell lines were generated by stimulating PBMCs isolated from two A/H5N1-naïve HLA-A*33:03-positive donors with each of the six peptide pools. On day 14, T cells were re-stimulated with peptide-loaded C1R.A*33:03 cells to measure CD8^+^ T cell activation via IFNγ production. For Donor 1, significantly robust CD8^+^IFNγ^+^ T cell responses were observed for pools 1, 3, 5 and 6 (**Figure 4A**), with Donor 2 showing strong reactivity towards pools 1, 5 and 6 (**Figure 4B**).

**Figure 4.**
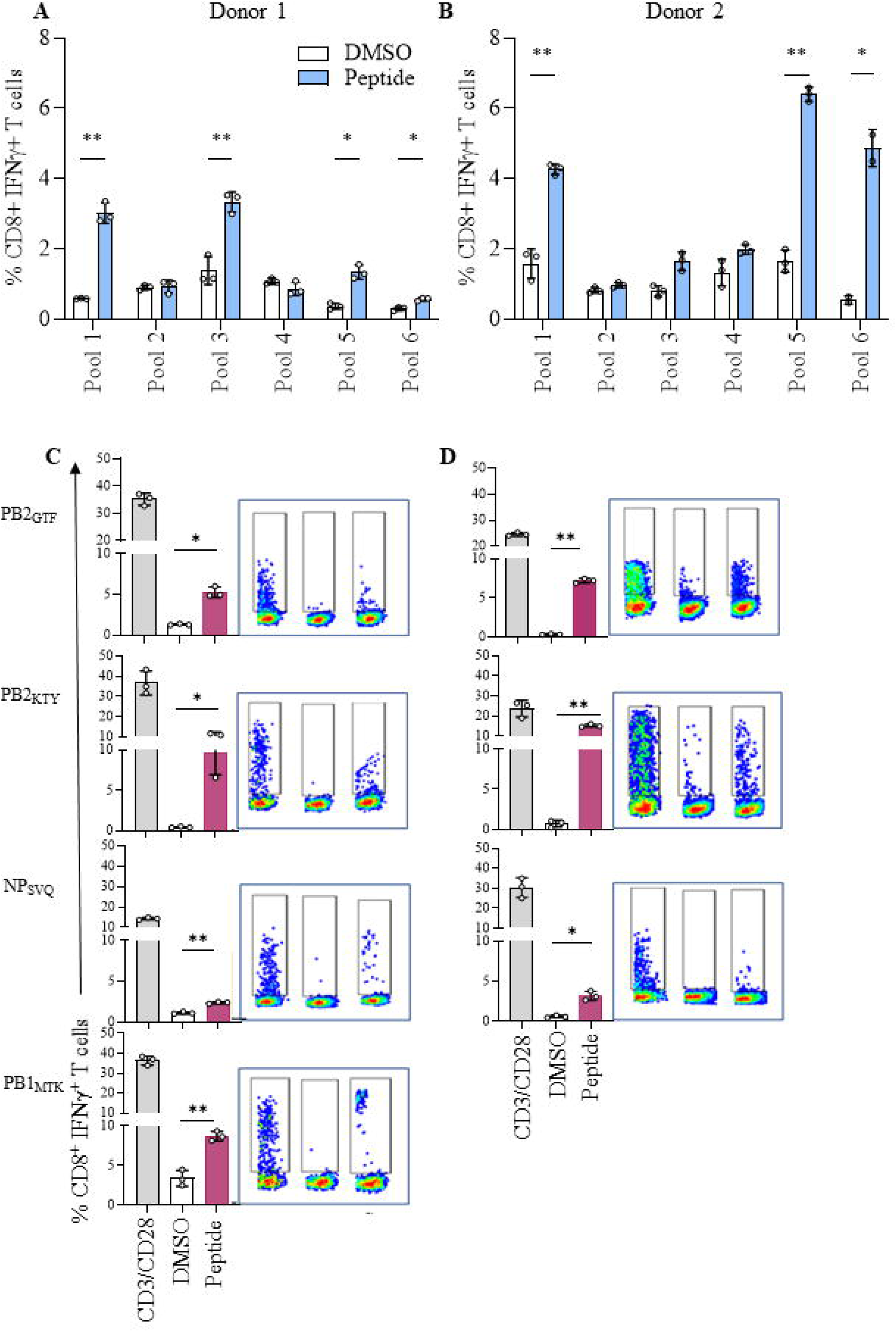
Identification of immunogenic A/H5N1- and A/X-31-specific CD8^+^ T cell epitopes. (A, B). Frequency of CD8^+^IFNγ^+^ T cells following stimulation with IAV (A/X-31 and A/H5N1) peptide pools in the presence of C1R.A*33:03 APCs. Experiments were conducted using one T-cell culture per condition, with peptide stimulation performed in triplicate as technical replicates. Statistical analysis was performed to identify significant differences between DMSO (peptide vehicle) and peptide loaded APC using a paired t test. A p value threshold was used to determine significance [*p<0.05, **p<0.01] (GraphPad Prism 9.0.0, USA). **(C, D)** Frequency of CD8^+^IFNγ^+^ T cells and corresponding concatenated dot plots showing individual peptide stimulation from pools 1 (PB2_GTF_ and PB1_MTK_, 3 (PB2_KTY_) and 5 (NP_SVQ_). PB1_MTK_ was oly tested in donor 1. Each assay was performed in technical triplicate. One-way ANOVA followed by Dunnett’s MCT was performed for data analysis [*p<0.05, **p<0.01].

To identify the immunogenic peptide within the positive response pools, PBMCs were stimulated with individual peptides. Here, day 14 CD8^+^IFNγ^+^ T cells from Donor 1 demonstrated significant recognition of three peptides (PB2_GTF_, PB2_KTY_, NP_SVQ_) from A/H5N1 and one peptide (PB1_MTK_) from A/X-31 (**Figure 4C**). For Donor 2, the same three A/H5N1 peptides displayed significant immunogenicity (**Figure 4D**). PB1_MTK_ peptide was not tested in Donor 2, and pool 6 peptides were not evaluated in either donor due to limited PBMC availability. As donors were A/H5N1-naïve, responses likely reflect cross-reactive memory T cells primed by seasonal influenza.

### Private dominant TCR clonotypes target the A/H5N1 NP_SVQ_ epitope

To examine the A/H1N1 epitope-specific αβTCR repertoire, day 14 NP_SVQ_-specific CD8^+^IFNγ^+^ T cells were single-cell sorted from each donor (**Figures 4C, D**). Characterisation of the NP_SVQ_-specific TCRs revealed dominant but distinct clonotypes in each donor, demonstrating private TCR usage, with Donor 1 expressing TRAV20.TRAJ13_TRBV27.TRBJ2-3 (paired αβTCR *Variable*_*Junction* genes; n=20/36 pairs, **Figure 5A**) and Donor 2 expressing TRAV14/DV4.TRAJ43_TRBV10-2.TRBJ2-1 (n=7/20; **Figure 5B**). Complete TCR repertoire information is reported in **Supplementary Data 3**.

**Figure 5.**
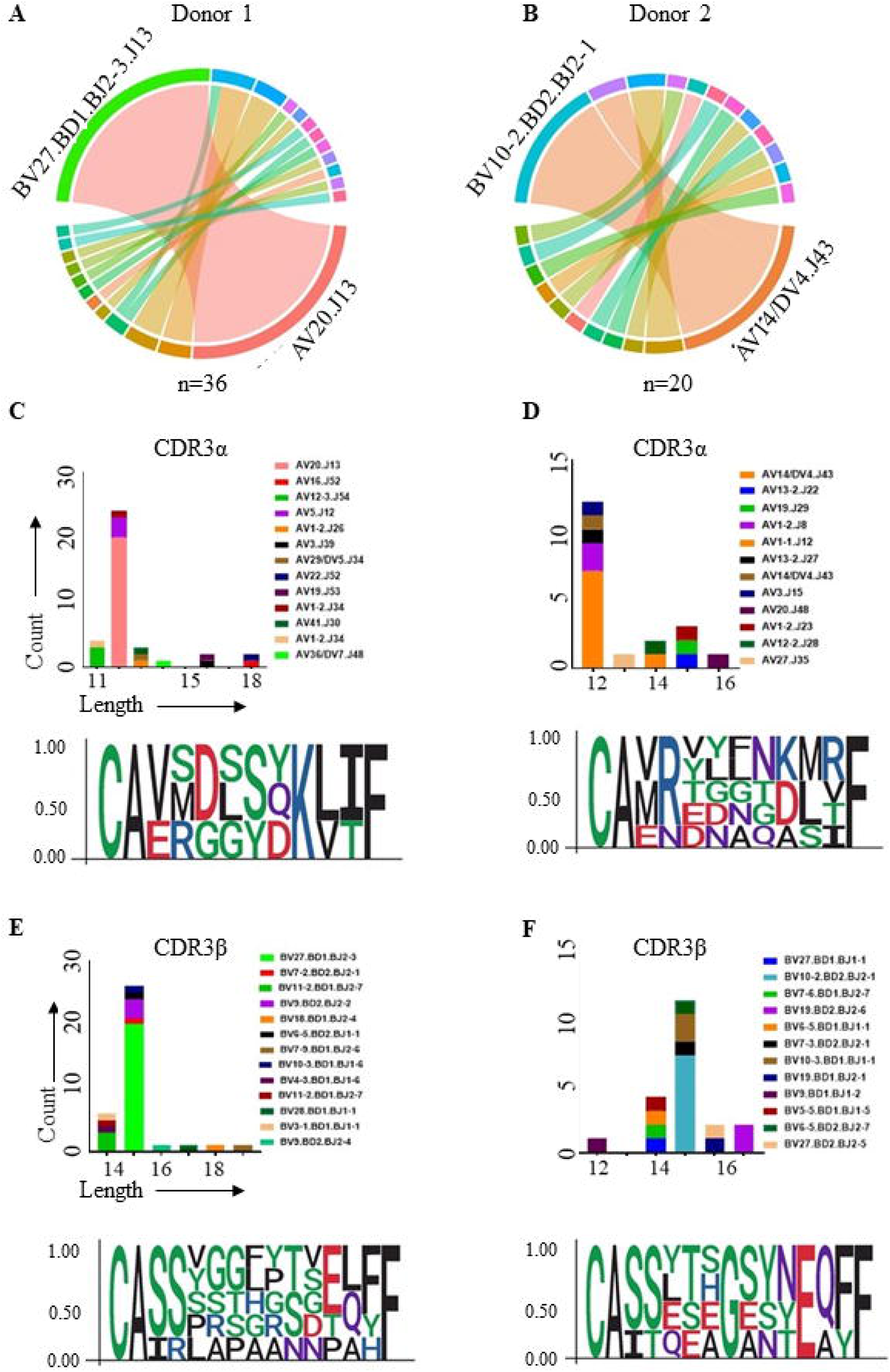
Private αβTCR clonotypes recognise the influenza A/H5N1 HLA-A*33:03/NP_SVQ_ epitope. (A, B). Chord plots of αβTCR repertoire usage for each donor recognising the NP_SVQ_ peptide. Connected ribbons represent paired α chain (TRAV/TRAJ) and β chain (TRBV/TRBD/TRBJ) usage. **(C, D)**. CDR3α and CDR3β length distributions. **(E, F).** Sequence logos showing amino acids residue variation in the CDR3 loops of dominant TRAV (13mer) and TRBV (15mer) genes. All sequencing analyses were performed using TCR_Explore [87].

The TCR CDR3α and CDR3β loops are important for direct interaction with the peptide presented by the HLA [45]. The CDR3α length distribution of both TRAV20 and TRAV14/DV4 encoded a 12mer in both donors, with dominant sequences of CAVSGGYQKVTF and CAMREYNNDMRF, respectively (**Figure 5C, D**). The CDR3β length distribution for both TRBV chains encoded a 15mer in both donors, with dominant sequences of CASSSGPHRTDTQYF and CASTQTSGSYNEQFF, respectively (**Figure 5E, F**). Collectively, these data demonstrated that the two dominant NP_SVQ_ TCR clonotypes were private, as they were not expressed by the alternate donor.

### High epitope conservation supports broad vaccine potential

Epitope conservation across divergent IAV strains is a key determinant of cross-protective T cell vaccine design. To evaluate the translational potential of the identified HLA-A33-restricted peptides, a conservation analyses was performed across available human and avian IAV sequences deposited in the NCBI Virus database [46].

Conservation of canonical A/X-31 peptides presented by HLA-A*33 allotypes were assessed in conjunction with predicted binding affinity (**Figure 6A, B**). Among these, 23 (34%) HLA-A*33:01-restricted and 20 (37%) HLA-A*33:03-restricted peptides were highly conserved (>90%) across circulating IAV strains. Importantly, most of these conserved peptides were predicted binders to their corresponding alleles (70% for HLA-A*33:01 and 60% for HLA-A*33:03) supporting their potential for stable presentation across diverse viral backgrounds. When extended to avian strains, approximately 70% of HLA-A*33:01 and HLA-A*33:03 peptides were conserved across A/H5N1 from avian and other hosts (**Figure 6C, D**), including the contemporary clade 2.3.4.4b viruses associated with recent outbreaks in humans and mammals [47].

**Figure 6.** Binding affinity and sequence conservation of identified A/X-31 and A/H5N1 peptides (A-D). Plots showing predicted binding affinity (%EL rank) versus sequence conservation of identified influenza A/X-31 virus peptides. Each dot represents an individual peptide and is colour-coded according to its source protein (HA, M1, NA, NP, NS1, NS2, PA, PB1, and PB1-F2), with the total number of analysed sequences for each protein indicated in parentheses in the legend. The y-axis represents predicted binding affinity expressed as %EL rank, where lower values indicate stronger predicted binding. The horizontal dotted line marks the threshold for predicted binders. The x-axis represents amino acid sequence conservation across circulating strains, and the vertical dotted line indicates highly conserved peptides (>90% conservation). Peptides located in the lower-right quadrant therefore represent strong predicted binders that are highly conserved. **E.** Conservation of the immunogenic peptides identified by CD8^+^ T cell assay in both human and avian hosts. PB1_MTK_ conservation has also been shown following interchange of residues from isoleucine to valine.

Importantly, two peptides, M1_FTL_ and PB2_DLN_ demonstrated >90% conservation across circulating human and avian A/H5N1 strains, were identified from both A/X-31 and A/H5N1 datasets, highlighting them as particularly promising vaccine candidates for individuals expressing these HLA allotypes. Among the four experimentally validated immunogenic epitopes (**Figure 4**), PB2_GTF_ exhibited the highest conservation across human and avian IAV strains and was present in both A/X-31 and the representative A/H5N1 strain used in this study (A/Chicken/Laos/Xaythiani-26/2006) (**Figure 6E**). In contrast, PB2_KTY_ and NP_SVQ_ showed higher conservation in avian relative to human isolates, whereas PB1_MTK_ displayed lower overall conservation. Notably, the previously reported I6V substitution within PB1_MTK_ is now fixed in the majority of circulating human and avian IAV strains [48], indicating evolutionary stabilisation of this variant.

Amino acid residue analysis across A/H3N2, A/H1N1, and A/H5N1 strains revealed limited amino acid variability (**Supplementary Figure 4**). The PB2_KTY_ epitope was >96.5% conserved in A/H5N1 and A/H1N1 strains, whereas 89% of A/H3N2 isolates carried an E5D substitution. Similarly, the NP_SVQ_ epitope was >95.4% conserved in A/H5N1 and A/H1N1 strains, while a conservative R10K substitution was observed in >73% of A/H3N2 isolates. In contrast, PB1_MTK_ showed greater similarity between A/H5N1 and A/H3N, whilst PB2_GTF_ remained >98.6% conserved across all viral strains.

Collectively, the high conservation and maintained predicted binding capacity of these CD8⁺ T cell epitopes support their potential utility in broadly protective T cell–based influenza vaccines, particularly in populations expressing HLA-A33.

## Discussion

The emergence of avian influenza A/H5N1 clade 2.3.4.4b in U.S. cattle, with documented human infections, reinforces the pandemic potential of this virus [49]. Current antibody-inducing vaccines provide limited cross-strain protection due to rapid antigenic drift and shift, highlighting the need for next generation strategies that elicit durable cellular immunity against influenza [50]. CD8⁺ T cells targeting conserved internal proteins represent a promising approach [51, 52]. However, HLA polymorphism remains a major challenge for broad vaccine design [53]. Identification of highly conserved, immunogenic epitopes presented by prevalent HLA allotypes is therefore essential, particularly for Asian populations where avian influenza A/H5N1 remains endemic [54].

We characterised the immunopeptidomes of HLA-A*33:03, predominant in South Asia, and the closely related HLA-A*33:01, common in East and Southeast Asia [37]. To our knowledge, this is the first study to define HLA-A*33:03-restricted ligands derived from avian A/H5N1 and assess their immunogenicity. We identified 100 novel A/X-31 peptides and 81 novel A/H5N1 peptides across both allotypes, substantially expanding the HLA-A33 influenza epitope repertoire.

Despite belonging to the HLA-A3 supertype, HLA-A*33:01 and HLA-A*33:03 displayed marked divergence in peptide presentation, with ∼70% non-overlapping viral repertoires. Notably, HLA-A*33:01 showed a subtle but consistent enrichment of aspartic acid (D) at the P1 position, likely due to the presence of histidine (H) at position 171 within the A pocket. Histidine, being positively charged at physiological pH, may facilitate stronger ionic interactions with the negatively charged side chain of aspartic acid. In contrast, HLA-A*33:03 contains a tyrosine (Y) at this position, which is neutral and lacks the same electrostatic potential, resulting in no marked preference between aspartic acid (D) and glutamic acid (E) at P1.

HLA-A*33:01 predominantly presented canonical 9mer peptides, whereas HLA-A*33:03 favored longer peptides and exhibited a higher proportion of predicted binders, suggesting qualitative differences in peptide selection. Although HLA class I molecules have historically been associated with 9mer ligands [55], structural and functional studies have demonstrated that peptides of variable length can be accommodated through conformational flexibility of the binding groove and peptide bulging, while still eliciting effective CD8⁺ T cell responses [56, 57]. The broader length repertoire observed for HLA-A*33:03 therefore suggests allele-specific structural constraints that may influence peptide selection and TCR engagement. Indeed, CD8⁺ T cell responses in individuals expressing these alleles may be directed toward largely different antigenic landscapes, potentially influencing immunodominance hierarchies and viral escape pathways.

Differences were also observed in predicted binding capacity, with a greater proportion of peptides classified as binders for HLA-A*33:03 than HLA-A*33:01 in both uninfected datasets (70% vs 48%) and A/X-31–derived peptides (74% vs 62%). This suggests distinct peptide stabilisation thresholds or structural properties of the binding groove may influence allele-specific epitope selection and downstream immune responsiveness [58].

Although HLA supertypes are defined based on shared primary anchor residue motifs [59], our data demonstrate that substantial divergence in peptide repertoires can occur within the same supertype. Even when identical amino acids are tolerated at the C-terminal anchor (PΩ), differences in residue preference and binding strength can markedly influence peptide selection [60]. Comparative analysis across three members of the HLA-A3 supertype (HLA-A*11:01, HLA-A*33:01, and HLA-A*33:03) revealed strikingly limited peptide overlap, underscoring the challenges of supertype-based epitope formulations.

Within the broader HLA-A3 supertype family, 56 IAV epitopes have been reported in IEDB to drive CD8⁺ T cell responses [58]. Our study expands this repertoire by identifying multiple HLA-A33-restricted epitopes, including four immunogenic HLA-A*33:03-restricted CD8⁺ T cell epitopes; two novel and two previously described for other HLA-A3 supertype alleles. Notably, three of these epitopes exhibit high conservation (>90%) across circulating human IAV strains, including clade 2.3.4.4b A/H5N1 viruses [61].

Strikingly, we detected A/H5N1-specific CD8⁺ T cell responses in two naïve HLA-A*33:03 positive individuals, suggesting that these responses likely from prior exposure to seasonal IAV that conferred cross-strain protection. Our findings align with previous reports showing that healthy individuals with seasonal influenza-specific CD4⁺ and CD8⁺ memory T cells exhibit broad cross-reactivity to conserved internal proteins, including NP and M1, from avian A/H5N1 viruses [62, 63]. Consistent with this, conservation analysis of the four immunogenic peptides derived from PB2, PB1, and NP demonstrated high sequence similarity across circulating A/H1N1, A/H3N2, and A/H5N1 strains, with predominantly single amino acid substitutions observed between subtypes. Here, two of the epitopes identified, NP_SVQ_ and PB2_GTF_, have also been reported as HLA-A3 supertype restricted epitopes [38, 64]. Despite limited overall overlap among A3 supertype alleles, the presence of these shared epitopes suggests that broader cross-presentation within the supertype is possible, supporting their potential inclusion in universal influenza vaccine strategies targeting the HLA-A3 supertype.

Current understanding of human IAV-specific αβTCR repertoires has largely centered on the HLA-A*02:01-restricted M1_58-66_ epitope [65–67], which exhibits a strong public bias toward paired TRAV27_TRBV19 clonotypes [68–70]. In contrast, HLA-A*03:01-restricted NP_265-273_ responses are more diverse and predominately private [71]. Although the HLA-A*33:03-restricted CD8^+^ T cell recognition of NP_SVQ_ has been described [64], its TCR architecture has not been defined. Here we demonstrate that NP_SVQ_-specific responses in two unrelated donors were dominated by distinct private clonotypes; TRAV20.TRAJ13_TRBV27.TRBJ2-3 (55.55%) for Donor 1 and TRAV14/DV4.TRAJ43_TRBV10-2.TRBJ2-1 (35%) for Donor 2. Despite differing gene usage, both repertoires shared features, including CDR3α (12mer) and CDR3β (15mer) lengths. These CDR3 regions are slightly longer than those reported for other immunodominant IAV epitopes (typically 7-10 amino acids) such as HLA-B*37:01-restricted NP_338-346_ [72], HLA-A*02:01-restricted M1_58-66_ [69] and HLA-B*35:01-restricted NP_418-426_ [69], suggesting allele-and epitope-specific structural constraints that shape TCR selection and antigen recognition.

Taken together, our findings demonstrate that despite minimal sequence divergence, HLA-A*33:01 and HLA-A*33:03 present largely distinct influenza-derived peptide repertoires with differing length preferences and predicted binding affinities. The broader representation of conserved peptides and higher proportion of predicted binders observed for HLA-A*33:03 suggest it may support qualitatively enhanced cellular immunity to influenza. Direct functional comparisons between the two allotypes were not feasible due to the absence of T cell response data for HLA-A*33:01 and warrant future investigation. These results underscore functional diversity even among closely related HLA allotypes and challenge assumptions of redundancy within supertypes. Incorporating newly defined, conserved HLA-A33-restricted epitopes into T cell-based universal vaccine startegies may therefor improve population coverage. Given the high prevalence of HLA-A*33:03 within South Asian populations, such allotype-informed strategies could be particularly valuable for enhancing vaccine coverage and protection in regions where avian influenza viruses are endemic.

## Materials and Methods

### Human samples

Human experimental work was conducted according to both the Declaration of Helsinki Principles and the Australian National Health and Medical Research Council (NHMRC) Code of Practice. Peripheral blood mononuclear cells (PBMCs) were isolated from blood using Ficoll density gradient centrifugation and cryopreserved as previously described [73]. The study was approved by Monash University Human Research Ethics Committee (Project ID 22209), with all blood donors providing written informed consent and recruitment between 7/10/2019-15/10/2024.

### Cell lines, influenza virus and peptides

The B-lymphoblastic cell line C1R (Class I-reduced) is a mutant antigen-presenting cell (APC) that does not express HLA-A but expresses low levels of HLA-B*35:03 and normal levels HLA-C*04:01 [74, 75]. C1R.A*33:03 and C1R.A*33:01 cells were generated by transfecting parental C1R cells with a construct encoding HLA-A*33:03 using the pcDNA3.1(+)-hygro backbone [76] or retroviral transduction of HLA-A*33:01 using a pMIG-GFP vector-based approach [77]. Cell lines were maintained in RF10 medium (RPMI-1640 supplemented with 10% heat-inactivated fetal bovine serum (FBS; Gibco, Thermo Fisher Scientific, USA), 100 mM MEM non-essential amino acids (Gibco), 55 mM 2-mercaptoethanol (Gibco), 5 mM HEPES buffer solution (Gibco), 1 mM MEM sodium pyruvate (Gibco), 1 mM L-glutamine (Gibco), 100 U/mL penicillin and 100 mg/mL streptomycin (Gibco)). C1R.A*33:03 required 0.3 mg/mL hygromycin B (Thermo Fisher Scientific) for HLA selection maintenance. Cell lines were tested [78] and shown to be negative for mycoplasma during all experimental procedures. Influenza A (A/X-31) virus (laboratory strain with seasonal H3N2 surface proteins and internal proteins from H1N1 PR8) was grown in the allantoic cavity of day 10–embryonated chicken eggs for 3 days at 35°C with viral titres determined by plaque assay on Madin-Darby canine kidney (MDCK) cells. Influenza peptides were synthesised by Mimotopes (Melbourne, Australia) using standard Fmoc chemistry and purified to a purity >90%. Peptides were reconstituted in dimethyl sulfoxide (DMSO; Sigma-Aldrich) at a final stock solution concentration 10 mM and aliquots stored at-80°C.

### *In vitro* infection of C1R cells with A/X-31 for immunopeptidomics

Cell lines (C1R, C1R.A*33:03 and C1R.A*33:01) were cultured in 1700 cm^2^ filter-capped roller bottles (Corning, USA) containing 800 mL of RF5 (same composition as RF10 except 5% FBS), slowly rotating at 37°C, 5% CO_2_ until 1×10^9^ cells were harvested. Cells were washed in phosphate buffer saline (PBS), centrifuged (1282 *g*, 15 min, room temperature [RT]) and reseeded in 50 mL tubes at a density of 1×10^7^ cells/mL in RPMI. Cells were infected with influenza A/X-31 virus at a Multiplicity of Infection (MOI) of 5 and the viral co-culture maintained at 37°C, 5% CO_2_ with slow rotation. After 1 hr, cells were returned to roller bottles with the addition 400 mL of 1:1 conditioned media (media from the harvested cells): 400 mL RF10 and incubated for a further 11 hrs (slow rotation at 37°C, 5% CO_2_). To validate HLA expression and IAV infection efficacy, an aliquot of 1×10^6^ cells were surface stained with anti-HLA-I PE-Cy7 (clone W6/32; BioLegend, USA) for 30 min at 4°C. Cells were washed in PBS and then fixed in 1% paraformaldehyde (ProSciTech, Australia) in PBS for 20 min at RT. Fixed cells were then permeabilised (0.3% saponin in PBS) and stained with anti-influenza A NP-FITC mAb (clone 1331; GeneTex, USA) for 30 min at 4°C. Cells were washed in PBS and both HLA and NP expression measured on a Becton Dickinson (BD) LSRII flow cytometer and the data analysed using FlowJo software (version 10, BD). Remaining influenza-infected cells were harvested into 500 mL V-bottom flasks (Corning), centrifuged (1282 *g*, 15 min, 4°C) and supernatant discarded. Cell pellets were washed in PBS, centrifuged (1282 *g*, 15 min, 4°C) and supernatant discarded before being snap frozen in liquid nitrogen. The same procedure was used to generate equivalent uninfected cell pellets. Cell pellets were stored at −80°C until use.

### Expression of A/H5N1 protein antigens in C1R cells

Four viral proteins (HA, PB2, NP and M1) from A/H5N1 viral strains (A/Chicken/Laos/Xaythiani-26/2006) were selected for investigation. Full length protein sequences were accessed from the Influenza Virus Resource, NCBI [79] and cDNA encoding those sequences synthesised individually as a gene block (gBlock; Integrated DNA Technologies, USA). gBlocks were cloned into the pIRES2 DsRed-Express2 vector (Takarabio, Japan) and electroporated into C1R.A*33:01 and C1R.A*33:03 cells. After electroporation, cells were grown under antibiotic selection (0.5 mg/ml, Geneticin, Merck) and checked for dual expression of GFP (reporter for HLA expression) and DsRed (reporter for influenza antigen expression) by flow cytometry as above. Transfected cells were then sorted based on high GFP and DsRed expression using a BD Influx flow cytometer. These transfected C1R.A*33:03 and C1R.A*33:01 cell lines expressing a single A/H5N1 protein (*i.e.* HA, NP, PB2 or M1) were cultured in 1700 cm^2^ filter-capped roller bottles (Corning) containing 800 mL of RF5 media, slowly rotating at 37°C, 5% CO_2_ until 1×10^9^ cells were harvested, washed in PBS, snap frozen and stored as described above.

### Immunopeptidomics and identification of HLA-bound peptides by mass spectrometry

Cell pellets were lysed using a combination of mechanical cryomilling (Retsch Mixer Mill MM 400) and detergent-based lysis (0.5% IGEPAL CA-630, 50 mM Tris-HCl pH 8.0, 150 mM NaCl and protease inhibitors [cOmplete Protease Inhibitor Cocktail Tablet; Roche Molecular Biochemicals]) for 1 hr at 4°C. Cell lysates were cleared by ultracentrifugation and solubilised HLA-I complexes isolated by immunoaffinity purification using protein-A agarose-bound pan HLA-I antibody (W6/32) as described previously [80]. Peptides were dissociated from the HLA-I complexes by treatment with 10% acetic acid and extracted peptides fractionated by offline reversed-phase high performance liquid chromatography (RP-HPLC) as described [81]. Peptide-containing fractions were combined into nine concatenated pools, concentrated using vacuum centrifugation (LABCONCO, USA), then reconstituted in a final volume of 12 μL of 0.1% formic acid (FA; Thermo Fisher Scientific). These pooled and concentrated samples were analysed by liquid chromatography tandem mass spectrometry (LC-MS/MS) on a Q-Exactive Plus Hybrid Quadrupole Orbitrap (Thermo Fisher Scientific) coupled to a Dionex UltiMate 3000 RSLC nano system (Thermo Fisher Scientific) using a data-dependent acquisition (DDA) strategy [38]. Data were analysed using PEAKS Xpro software (Bioinformatic Solutions Inc., Canada). Spectra were searched against a custom proteome database comprising the human reference proteome (UniProt/Swiss-Prot v2024) together with either the influenza A/X-31 proteome generated from six-frame translation of viral genome or selected protein (HA, M1, NP and PB2) sequences from A/H5N1 strain (A/Chicken/Laos/Xaythiani-26/2006). Sequence motifs were generated utilising human and influenza combined search peptides assigned at confidences greater than that required for a 1% false discovery rate (FDR) using Seq2logo2.0 [82]. Of note, for influenza peptide identification, a 5% FDR cutoff and a peptide length of 8-14mer were applied to balance stringency and maximise the number of potential influenza-derived peptides for subsequent validation and functional testing. To minimise false influenza peptide identification, all influenza (A/X-31/A/H5N1)-derived peptides were manually checked against the corresponding uninfected datasets from C1R, C1R.A*33:01 and C1R.A*33:03 cells to show their likely viral origin. Peptide-binding affinities were predicted using antigen presentation mode of NetMHCpan-4.2 [44], with % EL rank determining strong binders (<0.5 SB), weak binders (<2 WB) and non-binders (>2 NB).

### Conservation analysis

Complete influenza A protein isolates and associated metadata were downloaded from NCBI Virus (captured 15/02/2026) [46]. Avaiable number of different influenza proteins sequences (mentioned in the figure) were then aligned with high fidelity using MAFFT G-INS-I algorithm [83] via M3 HPC (High-Performance Computing (HPC) cluster (eResearch, Monash University). The complete alignment was then filtered by the metadata (*e.g.,* genotype) for downstream analysis.

### *In vitro* expansion of IAV-specific CD8^+^ T cells

A total of 5 × 10^6^ HLA-A*33:03-positive cryopreserved PBMCs were thawed and washed twice in RPMI, with antigen-specific expansions performed as previously described [39]. Briefly, one-third of the PBMCs were pulsed with 1 μM of each peptide, either individually or as pooled influenza peptides (**Table 1**) for 1 hr at 37°C, 5% CO_2_, then washed twice in RPMI. Co-cultures of peptide-stimulated PBMC with autologous non-stimulated PBMCs were maintained in RH10 (same constituents as RF10, except 10% heat-inactivated human blood group AB serum substituted for FBS; Sigma-Aldrich) and incubated at 37°C, 5% CO_2_. From day 4 to 14, T cell cultures were supplemented with 20 U/mL of recombinant human interleukin-2 (IL-2; Peprotech, USA). On day 14, expanded T cells were harvested. A total of 5 × 10^6^ cells were cryopreserved for single-cell αβ T cell receptor (TCR) sequencing, while the remainder was used for intracellular cytokine staining (ICS) assays.

### T cell restimulation and intracellular cytokine staining

On day14, *in vitro* expanded T cells were harvested and restimulated at a 2:1 ratio with 1 μM of peptide-loaded APCs and incubated at 37°C, 5% CO_2_. Media and Dynabeads® Human T-Activator CD3/CD28 beads (Thermo Fisher Scientific) represented background and positive controls, respectively. After 2 hrs, 10 μg/mL of Brefeldin A (Sigma-Aldrich) was added. At 6 hrs, cells were pelleted and cell surface markers labelled with antibodies specific for CD4 PE (clone RPA-T4), CD8 PerCP-Cy5.5 (clone SK1) and LIVE/DEAD® fixable Aqua stain (Thermo Fisher Scientific) for 30 min at 4°C in the dark. Cells were washed in PBS and fixed with 1% paraformaldehyde (ProSciTech, Australia) in PBS for 20 min at RT in the dark. Cells were washed in PBS and permeabilised with 0.3% Saponin (Sigma-Aldrich) in PBS containing anti-IFNγ PE-Cy7 (clone B27) and anti-TNFα V450 (clone MAb11) antibodies and incubated overnight at 4°C in the dark. Cells were washed in PBS prior to flow cytometry. All antibodies were purchased from BD Pharmingen (USA) and titrated for optimal staining efficiency. A maximum of 50,000 lymphocytes were analysed on a BD Fortessa flow cytometer and the data analysed using FlowJo software (version 10, BD).

### Single-cell sequencing of A/H5N1-specific CD8^+^ T cells

Cryopreserved day 14 T cell lines were thawed and rested overnight in RH10. Up to 5 × 10^6^ T cells were co-cultured with either C1R.A*33:03 cells alone or in addition with 1 μM of NP 10mer peptide SVQRNLPFER (NP_SVQ_) at a 2:1 effector-to-target ratio in RH5 media. Co-cultures were maintained for 4 hrs at 37°C, 5% CO_2_. After incubation, cells were washed using cold wash buffer (0.5% FBS, 2 mM EDTA pH 8.0 in PBS), centrifuged at 285 *g* for 5 min at 4°C, and the supernatant was discarded. The IFNγ catch reagent antibody (Miltenyi Biotech) was added, cells were incubated on ice for 5 min, resuspended in 10 mL of pre-warmed RH5 media with 1 μM NP_SVQ_ peptide re-added and incubated for 45 min at 37°C with gentle rotation. Cells were washed, centrifuged (285 *g*, 5 min, 4°C) and the pellet was stained with allophycocyanin-conjugated IFNγ detection reagent (Miltenyi Biotech) and anti-CD8 FITC (clone HIT8a, BD) antibody for 20 min on ice. Cells were washed and centrifuged (285 *g*, 5 min, 4°C) with the pellet resuspended in 300 μL of cold wash buffer.

Single-cell sorting was conducted on a BD Influx flow cytometer, targeting CD8⁺IFNγ⁺ cells as the peptide-responsive population. Single-cell sorts were deposited directly into 96-well PCR plates (Bio-Rad, USA) and immediately stored at −80°C. Paired TCRα and TCRβ genes were amplified via multiplex nested RT-PCR using external and internal rounds of PCR that included 40 TRAV and 27 TRBV forward primers, and a TRAC and TRBC reverse primer [84]. Samples were Sanger sequenced at Micromon Genomics (Monash University) and sequences were analysed according to the ImMunoGeneTics/V-QUEry and STandardization web-based tool [85]. All TCR nomenclature was according to Folch *et al.* [86]. CDR3 amino acid sequences described within the text start from CDR3-position 3, which is equivalent to amino acid position 107 of the TRAV and TRBV segments, and end at TRAJ-position 10 or TRBJ-position 6. TCR_Explore was used to visualise the paired α- and β-chains, as well as to perform CDR3 length and motif analysis [87].

## Statistical analysis

All data were reported as mean ± standard deviation of the mean (SD), unless stated otherwise. Non-parametric unpaired data was analysed using Two-way ANOVA followed by Sidak’s multiple comparisons tests (MCT) or One-way ANOVA with Dunnett’s MCT, whereas paired data was analysed using Wilcoxon matched-pairs signed-rank test (GraphPad Prism 9.0.0, USA). Statistical significance is indicated by *p < 0.05, **p < 0.01, ***p<0.001 or ****p < 0.0001.

## Data availability

Mass spectrometry proteomics data and PEAKS online® search results, have been deposited to the ProteomeXchange Consortium via the PRIDE [88] partner repository with the dataset identifier PXD078870 and 10.6019/PXD078870 [89].

## Acknowledgements

The authors acknowledge Monash University platform facilities, FlowCore for flow cytometry and Micromon Genomics for Sanger sequencing and their technical support. Computational resources for proteomics analysis were supported by the R@CMon/Monash Node of the NeCTAR Research Cloud, an initiative of the Australian Government’s Super Science Scheme and the Education Investment Fund. The authors also acknowledge the provision of instrumentation, training and technical support by the Monash Proteomics and Metabolomics Platform. Computational resources for conservation analysis were supported by Monash eResearch capabilities, including M3 HPC. The authors are grateful to Steve Rockman, CSL Seqirus Ltd, Parkville, VIC, Australia for the laboratory strain of influenza A/X-31 (H3N2).

## Author Contributions

AKMM, PTI, NAM designed experiments. AKM, PTI, SW, PS, NAM performed experiments. MLB, AWP provided funding, reagents and intellectual input. AKMM, PTI, SW, PS, NAM, MJ, NPC analysed data. KK provided reagents. AKMM and NAM wrote the manuscript. All authors read and approved the manuscript.

## Conflict of Interest

AWP is a scientific advisor for Bioinformatics Solutions Inc (Canada), a shareholder and scientific advisor for Evaxion Biotech (Denmark), and a co-founder of Resseptor Therapeutics (Australia). These organisations had no role in the design of the study; in the collection, analyses, or interpretation of data; in the writing of the manuscript; or in the decision to publish the results.

All other authors declare that the research was conducted in the absence of any commercial or financial relationships that could be construed as a potential conflict of interest.

## Funding

This work was supported by the National Health and Medical Research Council of Australia (NHMRC) (Grants APP1122099 and APP2016596 to A.W.P.).

This study was conducted as part of a PhD project for AKMM, funded by Monash University through a co-funded Monash Graduate Scholarship.

## Supplementary Information

**Supplementary Figure 1. Positional amino acid usage of MS-identified 9-mer peptides presented by HLA-A*33:01 and HLA-A*33:03.**

Scatter plots (P1–P9) compare the amino acid frequency (%) at each position of 9-mer peptides identified by mass spectrometry and assigned to HLA-A*33:01 (x-axis) and HLA-A*33:03 (y-axis). Each circle represents a single amino acid at the indicated position. The dashed diagonal line indicates equal frequency between the two alleles. Amino acids highlighted in red denote residues exhibiting a ≥5% difference in positional frequency between HLA-A*33:01 and HLA-A*33:03. Aspartic acid (D) is strongly enriched in HLA-A*33:01-restricted peptides, whereas glutamic acid (E) is preferentially enriched in HLA-A*33:03-restricted peptides. Overall, the two alleles show highly similar amino acid distributions across most positions, with selective differences at specific sites.

**Supplementary Figure 2. Binding motifs HLA-A33-restricted 10-12mer peptides.**

**(Ai-iii).** A*33:01-bound longer peptides (10-12mer) showed similar motifs to 9mer peptides. **(Bi-iii).** A*33:03-bound longer peptides (10-12mer) showed similar motifs to 9mer peptides. Seq2Logo were used to generate sequence motifs (https://services.healthtech.dtu.dk/services/Seq2Logo-2.0/).

**Supplementary Figure 3. 9mer peptide binding motifs in A/H5N1 protein transfected cell lines.**

**A.** Binding motifs of 9mer peptides from all A/H5N1 transfected cell lines show similarities among themselves and also with those of naturally infected cells. **B**. 50% of A*33:03 restricted A/H5N1 peptides were predicted binders compared to 46% of the identified A/H5N1 peptides were binders to A*33:01. Peptide-binding affinities were predicted using NetMHCpan-4.2 [44], with % rank determining strong binders (<0.5 SB), weak binders (<2 WB), non-binders (>2 NB).

**Supplementary Figure 4: Residue level conservation of immunogenic peptides in different influenza strains.**

Sequence conservation analysis of four immunogenic peptides derived from PB2, PB1, and NP proteins across A/H1N1, A/H3N2, and A/H5N1 viruses. The consensus (most frequent) sequence for each peptide is shown at the top of each panel (blue boxes), with the percentage frequency and number of identical sequences indicated on the right. Variant sequences are listed below, with amino acid substitutions highlighted in red and their corresponding frequencies shown. The total number of sequences analysed for each strain is indicated in parentheses. Most peptides demonstrate high conservation (>95%) across circulating strains, with substitutions occurring at low frequencies, supporting their potential contribution to cross-strain CD8⁺ T cell immunity.

**Supplimentary Data 1: HLA-A-33-restricted influenza A/X-31 peptides**

**Supplimentary Data 2: HLA-A*33:01-bound peptides from all A/H5N1 transfected cells**

## Notes

### Summary of Updates

Supp Table 3 added as was originally omitted from submission

## References

1. Hsieh, Y.C., et al., Influenza pandemics: past, present and future. J Formos Med Assoc, 2006. 105(1): p. 1–6.

2. Influenza Virus. Transfus Med Hemother, 2009. 36(1): p. 32–39.

3. Barr, I.G. and K. Subbarao, Implications of the apparent extinction of B/Yamagata-lineage human influenza viruses. npj Vaccines, 2024. 9(1): p. 219.

4. WHO, Influenza at the human-animal interface. 2018.

5. Hurtado, T.R., Human Influenza A (H5N1): A Brief Review and Recommendations for Travelers. Wilderness & Environmental Medicine, 2006. 17(4): p. 276–281.

6. Verhagen, J.H., et al., Host Range of Influenza A Virus H1 to H16 in Eurasian Ducks Based on Tissue and Receptor Binding Studies. J Virol, 2021. 95(6).

7. Robert G. Webster, A.S.M., Thomas J. Braciale, and R.A. Lamb., Textbook of influenza. 2nd ed. 2013, West Sussex, UK: John Wiley & Sons, Ltd.

8. WHO, Avian Influenza Weekly Update # 1001: 13 June 2025. 2025, World Health Organization: Geneva, Switzerland. p. 4.

9. Nypaver, C., C. Dehlinger, and C. Carter, Influenza and Influenza Vaccine: A Review. J Midwifery Womens Health, 2021. 66(1): p. 45–53.

10. Koutsakos, M., et al., Human CD8(+) T cell cross-reactivity across influenza A, B and C viruses. Nat Immunol, 2019. 20(5): p. 613–625.

11. Hayward, A.C., et al., Natural T Cell-mediated Protection against Seasonal and Pandemic Influenza. Results of the Flu Watch Cohort Study. Am J Respir Crit Care Med, 2015. 191(12): p. 1422–31.

12. Clemens, E.B., et al., Towards identification of immune and genetic correlates of severe influenza disease in Indigenous Australians. Immunol Cell Biol, 2016. 94(4): p. 367–77.

13. Sridhar, S., et al., Cellular immune correlates of protection against symptomatic pandemic influenza. Nat Med, 2013. 19(10): p. 1305–12.

14. Quiñones-Parra, S.M., et al., A Role of Influenza Virus Exposure History in Determining Pandemic Susceptibility and CD8+ T Cell Responses. J Virol, 2016. 90(15): p. 6936–6947.

15. Grant, E.J., et al., Lack of Heterologous Cross-reactivity toward HLA-A*02:01 Restricted Viral Epitopes Is Underpinned by Distinct αβT Cell Receptor Signatures. J Biol Chem, 2016. 291(47): p. 24335–24351.

16. Valkenburg, S.A., et al., Protective efficacy of cross-reactive CD8+ T cells recognising mutant viral epitopes depends on peptide-MHC-I structural interactions and T cell activation threshold. PLoS Pathog, 2010. 6(8): p. e1001039.

17. Regner, M., et al., Cutting edge: rapid and efficient in vivo cytotoxicity by cytotoxic T cells is independent of granzymes A and B. The Journal of Immunology, 2009. 183(1): p. 37–40.

18. Metkar, S.S., et al., Human and mouse granzyme A induce a proinflammatory cytokine response. Immunity, 2008. 29(5): p. 720–33.

19. Thibault, P. and C. Perreault, Immunopeptidomics: Reading the Immune Signal That Defines Self From Nonself. Mol Cell Proteomics, 2022. 21(6): p. 100234.

20. Kulski, J.K., S. Suzuki, and T. Shiina, Human leukocyte antigen super-locus: nexus of genomic supergenes, SNPs, indels, transcripts, and haplotypes. Human Genome Variation, 2022. 9(1): p. 49.

21. Crux, N.B. and S. Elahi, Human Leukocyte Antigen (HLA) and Immune Regulation: How Do Classical and Non-Classical HLA Alleles Modulate Immune Response to Human Immunodeficiency Virus and Hepatitis C Virus Infections? Front Immunol, 2017. 8: p. 832.

22. Arora, J., et al., HLA Heterozygote Advantage against HIV-1 Is Driven by Quantitative and Qualitative Differences in HLA Allele-Specific Peptide Presentation. Mol Biol Evol, 2020. 37(3): p. 639–650.

23. Karnaukhov, V., et al., HLA variants have different preferences to present proteins with specific molecular functions which are complemented in frequent haplotypes. Frontiers in Immunology, 2022. Volume 13 - 2022.

24. Trolle, T., et al., The Length Distribution of Class I-Restricted T Cell Epitopes Is Determined by Both Peptide Supply and MHC Allele-Specific Binding Preference. J Immunol, 2016. 196(4): p. 1480–7.

25. Sant, S., et al., HLA-B*27:05 alters immunodominance hierarchy of universal influenza-specific CD8+ T cells. PLOS Pathogens, 2020. 16(8): p. e1008714.

26. Subbarao, K., et al., Characterization of an avian influenza A (H5N1) virus isolated from a child with a fatal respiratory illness. Science, 1998. 279(5349): p. 393–6.

27. Duan, L., et al., *The development and genetic diversity of H5N1 influenza virus in China*, *1996-2006*. Virology, 2008. 380(2): p. 243–54.

28. WHO. Cumulative number of confirmed human cases for avian influenza A(H5N1) reported to WHO, 2003-2019. 2019 09/04/2019 [cited 2019 27/05/2019]; Available from: https://www.who.int/influenza/human_animal_interface/2019_04_09_tableH5N1.pdf?ua=1.

29. Gutiérrez, R.A., et al., A(H5N1) Virus Evolution in South East Asia. Viruses, 2009. 1(3): p. 335–61.

30. Chowdhury, S., et al., The Pattern of Highly Pathogenic Avian Influenza H5N1 Outbreaks in South Asia. Trop Med Infect Dis, 2019. 4(4).

31. Salaheldin, A.H., et al., Potential Biological and Climatic Factors That Influence the Incidence and Persistence of Highly Pathogenic H5N1 Avian Influenza Virus in Egypt. Front Microbiol, 2018. 9: p. 528.

32. Fang, L.-Q., et al., Environmental Factors Contributing to the Spread of H5N1 Avian Influenza in Mainland China. PLOS ONE, 2008. 3(5): p. e2268.

33. Boon, A.C., et al., Host genetic variation affects resistance to infection with a highly pathogenic H5N1 influenza A virus in mice. J Virol, 2009. 83(20): p. 10417–26.

34. Ahmed, R., M.B. Oldstone, and P. Palese, Protective immunity and susceptibility to infectious diseases: lessons from the 1918 influenza pandemic. Nat Immunol, 2007. 8(11): p. 1188–93.

35. Ruiz-Hernandez, R., et al., Host genetics determine susceptibility to avian influenza infection and transmission dynamics. Sci Rep, 2016. 6: p. 26787.

36. Kim, Y., et al., Prevalence of Avian Influenza A(H5) and A(H9) Viruses in Live Bird Markets, Bangladesh. Emerg Infect Dis, 2018. 24(12): p. 2309–2316.

37. Gonzalez-Galarza, F.F., et al., Allele frequency net database (AFND) 2020 update: gold-standard data classification, open access genotype data and new query tools. Nucleic Acids Res, 2020. 48(D1): p. D783–D788.

38. Habel, J.R., et al., HLA-A*11:01-restricted CD8+ T cell immunity against influenza A and influenza B viruses in Indigenous and non-Indigenous people. PLoS Pathog, 2022. 18(3): p. e1010337.

39. Hensen, L., et al., CD8(+) T cell landscape in Indigenous and non-Indigenous people restricted by influenza mortality-associated HLA-A*24:02 allomorph. Nat Commun, 2021. 12(1): p. 2931.

40. Vita, R., et al., The Immune Epitope Database (IEDB): 2018 update. Nucleic Acids Res, 2019. 47(D1): p. D339–D343.

41. Wu, C., et al., Systematic identification of immunodominant CD8^+^ T-cell responses to influenza A virus in HLA-A2 individuals. Proceedings of the National Academy of Sciences, 2011. 108(22): p. 9178–9183.

42. Knight, M., et al., Imprinting, immunodominance, and other impediments to generating broad influenza immunity. Immunological Reviews, 2020. 296(1): p. 191–204.

43. van de Sandt, C.E., et al., Challenging immunodominance of influenza-specific CD8+ T cell responses restricted by the risk-associated HLA-A*68:01 allomorph. Nature Communications, 2019. 10(1): p. 5579.

44. Nilsson, J.B., et al., NetMHCpan-4.2: improved prediction of CD8+ epitopes by use of transfer learning and structural features. Frontiers in Immunology, 2025. Volume 16 - 2025.

45. Szeto, C., et al., TCR Recognition of Peptide-MHC-I: Rule Makers and Breakers. Int J Mol Sci, 2020. 22(1).

46. (MD), N.V.I.B., National Library of Medicine (US), National Center for Biotechnology Information.

47. Burrough, E.R., et al., *Highly Pathogenic Avian Influenza A(H5N1) Clade 2.3.4.4b Virus Infection in Domestic Dairy Cattle and Cats, United States*, *2024*. Emerg Infect Dis, 2024. 30(7): p. 1335–1343.

48. Squires, R.B., et al., Influenza Research Database: an integrated bioinformatics resource for influenza research and surveillance. Influenza and Other Respiratory Viruses, 2012. 6(6): p. 404–416.

49. Oguzie, J.U., et al., Avian Influenza A(H5N1) Virus among Dairy Cattle, Texas, USA. Emerg Infect Dis, 2024. 30(7): p. 1425–1429.

50. Chen, J. and Y.M. Deng, Influenza virus antigenic variation, host antibody production and new approach to control epidemics. Virol J, 2009. 6: p. 30.

51. Sridhar, S., et al., Cellular immune correlates of protection against symptomatic pandemic influenza. Nature medicine, 2013. 19(10): p. 1305–1312.

52. Thomas, P.G., et al., Cell-mediated protection in influenza infection. Emerg Infect Dis, 2006. 12(1): p. 48–54.

53. Daniels, M.A. and S.C. Jameson, Critical role for CD8 in T cell receptor binding and activation by peptide/major histocompatibility complex multimers. J Exp Med, 2000. 191(2): p. 335–46.

54. CDC, U., Highly Pathogenic Avian Influenza A(H5N1) in Birds and Other Animals. 2015, Centre for Disease Control, USA: Atlanta, USA.

55. Chen, Y., et al., Naturally processed peptides longer than nine amino acid residues bind to the class I MHC molecule HLA-A2.1 with high affinity and in different conformations. J Immunol, 1994. 152(6): p. 2874–81.

56. Burrows, J.M., et al., Preferential binding of unusually long peptides to MHC class I and its influence on the selection of target peptides for T cell recognition. Molecular Immunology, 2008. 45(6): p. 1818–1824.

57. Rist, M.J., et al., HLA peptide length preferences control CD8+ T cell responses. J Immunol, 2013. 191(2): p. 561–71.

58. Paul, S., et al., HLA class I alleles are associated with peptide-binding repertoires of different size, affinity, and immunogenicity. J Immunol, 2013. 191(12): p. 5831–9.

59. Sidney, J., et al., HLA class I supertypes: a revised and updated classification. BMC Immunol, 2008. 9: p. 1.

60. Desmet, J., et al., Anchor profiles of HLA-specific peptides: Analysis by a novel affinity scoring method and experimental validation. Proteins: Structure, Function, and Bioinformatics, 2005. 58(1): p. 53–69.

61. Xie, Z., et al., Clade 2.3.4.4b highly pathogenic avian influenza H5N1 viruses: knowns, unknowns, and challenges. Journal of Virology, 2025. 99(6): p. e00424–25.

62. Sidney, J., et al., Targets of influenza Human T cell response are mostly conserved in H5N1. bioRxiv, 2024: p. 2024.09.09.612060.

63. Lee, L.Y., et al., Memory T cells established by seasonal human influenza A infection cross-react with avian influenza A (H5N1) in healthy individuals. J Clin Invest, 2008. 118(10): p. 3478–90.

64. Alexander, J., et al., Identification of broad binding class I HLA supertype epitopes to provide universal coverage of influenza A virus. Hum Immunol, 2010. 71(5): p. 468–74.

65. Wang, Z., et al., Clonally diverse CD38(+)HLA-DR(+)CD8(+) T cells persist during fatal H7N9 disease. Nat Commun, 2018. 9(1): p. 824.

66. Glanville, J., et al., Identifying specificity groups in the T cell receptor repertoire. Nature, 2017. 547(7661): p. 94–98.

67. Nguyen, T.H.O., et al., Perturbed CD8(+) T cell immunity across universal influenza epitopes in the elderly. J Leukoc Biol, 2018. 103(2): p. 321–339.

68. Gil, A., et al., Narrowing of human influenza A virus-specific T cell receptor α and β repertoires with increasing age. J Virol, 2015. 89(8): p. 4102–16.

69. Valkenburg, S.A., et al., Molecular basis for universal HLA-A*0201-restricted CD8+ T-cell immunity against influenza viruses. Proc Natl Acad Sci U S A, 2016. 113(16): p. 4440–5.

70. Chen, G., et al., Sequence and Structural Analyses Reveal Distinct and Highly Diverse Human CD8(+) TCR Repertoires to Immunodominant Viral Antigens. Cell Rep, 2017. 19(3): p. 569–583.

71. Nguyen, A.T., et al., Homologous peptides derived from influenza A, B and C viruses induce variable CD8(+) T cell responses with cross-reactive potential. Clin Transl Immunology, 2022. 11(10): p. e1422.

72. Grant, E.J., et al., Broad CD8(+) T cell cross-recognition of distinct influenza A strains in humans. Nat Commun, 2018. 9(1): p. 5427.

73. Menon, T., et al., CD8(+) T-cell responses towards conserved influenza B virus epitopes across anatomical sites and age. Nat Commun, 2024. 15(1): p. 3387.

74. Edwards, P.A., et al., A human-hybridoma system based on a fast-growing mutant of the ARH-77 plasma cell leukemia-derived line. Eur J Immunol, 1982. 12(8): p. 641–8.

75. Zemmour, J., et al., The HLA-A,B “negative” mutant cell line C1R expresses a novel HLA-B35 allele, which also has a point mutation in the translation initiation codon. J Immunol, 1992. 148(6): p. 1941–8.

76. Braun, A., et al., Mapping the immunopeptidome of seven SARS-CoV-2 antigens across common HLA haplotypes. Nat Commun, 2024. 15(1): p. 7547.

77. Alekseeva, N.A., et al., Obtaining Gene-Modified HLA-E-Expressing Feeder Cells for Stimulation of Natural Killer Cells. Pharmaceutics, 2024. 16(1).

78. Siegl, D., et al., A PCR protocol to establish standards for routine mycoplasma testing that by design detects over ninety percent of all known mycoplasma species. iScience, 2023. 26(5): p. 106724.

79. Virus, N. [cited 2025 17 January]; Available from: https://www.ncbi.nlm.nih.gov/labs/virus/vssi/#/.

80. Purcell, A.W., S.H. Ramarathinam, and N. Ternette, Mass spectrometry-based identification of MHC-bound peptides for immunopeptidomics. Nat Protoc, 2019. 14(6): p. 1687–1707.

81. Thomson, P.J., et al., Modification of the cyclopropyl moiety of abacavir provides insight into the structure activity relationship between HLA-B*57:01 binding and T-cell activation. Allergy, 2020. 75(3): p. 636–647.

82. Thomsen, M.C. and M. Nielsen, Seq2Logo: a method for construction and visualization of amino acid binding motifs and sequence profiles including sequence weighting, pseudo counts and two-sided representation of amino acid enrichment and depletion. Nucleic Acids Res, 2012. 40(Web Server issue): p. W281-7.

83. Katoh, K. and D.M. Standley, MAFFT Multiple Sequence Alignment Software Version 7: Improvements in Performance and Usability. Molecular Biology and Evolution, 2013. 30(4): p. 772–780.

84. Wang, G.C., et al., T cell receptor αβ diversity inversely correlates with pathogen-specific antibody levels in human cytomegalovirus infection. Sci Transl Med, 2012. 4(128): p. 128ra42.

85. Balakrishnan, A. and G.P. Morris, The highly alloreactive nature of dual TCR T cells. Curr Opin Organ Transplant, 2016. 21(1): p. 22–8.

86. Ferrara, G.B., et al., Bone marrow transplantation from unrelated donors: the impact of mismatches with substitutions at position 116 of the human leukocyte antigen class I heavy chain. Blood, 2001. 98(10): p. 3150–5.

87. Mullan, K.A., et al., TCR_Explore: A novel webtool for T cell receptor repertoire analysis. Comput Struct Biotechnol J, 2023. 21: p. 1272–1282.

88. Perez-Riverol, Y., et al., The PRIDE database at 20 years: 2025 update. Nucleic Acids Res, 2025. 53(D1): p. D543–d553.

89. Deutsch, E.W., et al., The ProteomeXchange consortium at 10 years: 2023 update. Nucleic Acids Research, 2023. 51(D1): p. D1539–D1548.

