## Supplementary figures and images for "Identification of HLA-A33-restricted CD8^+^ T cell epitopes from avian influenza A/H5N1"

### Figure S1

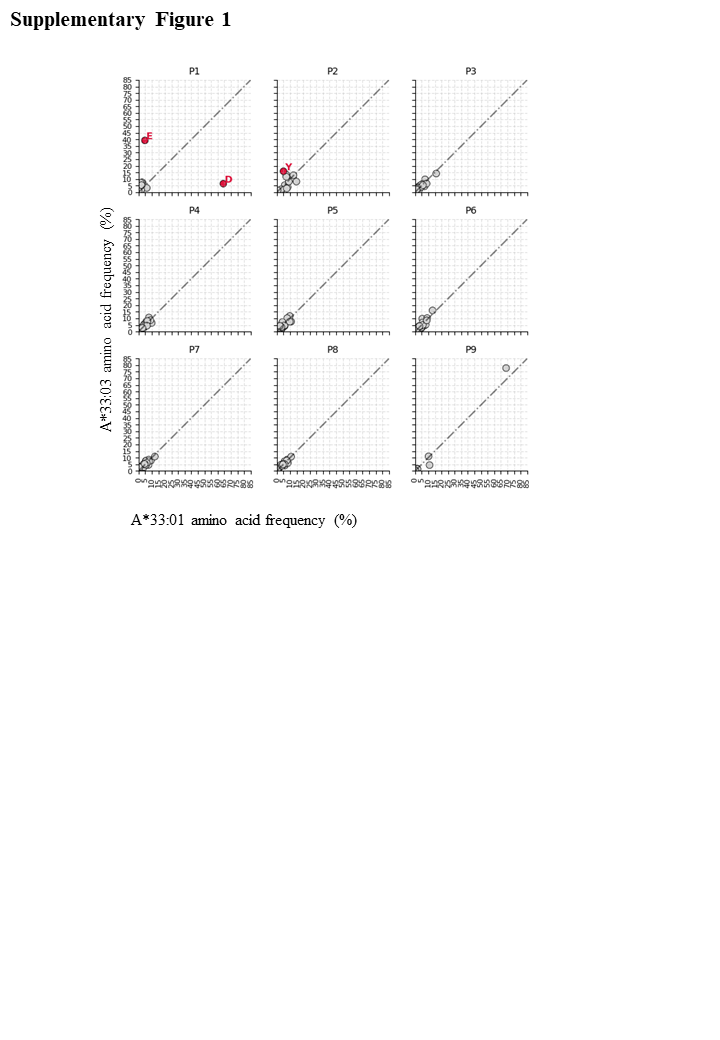

### Figure S2

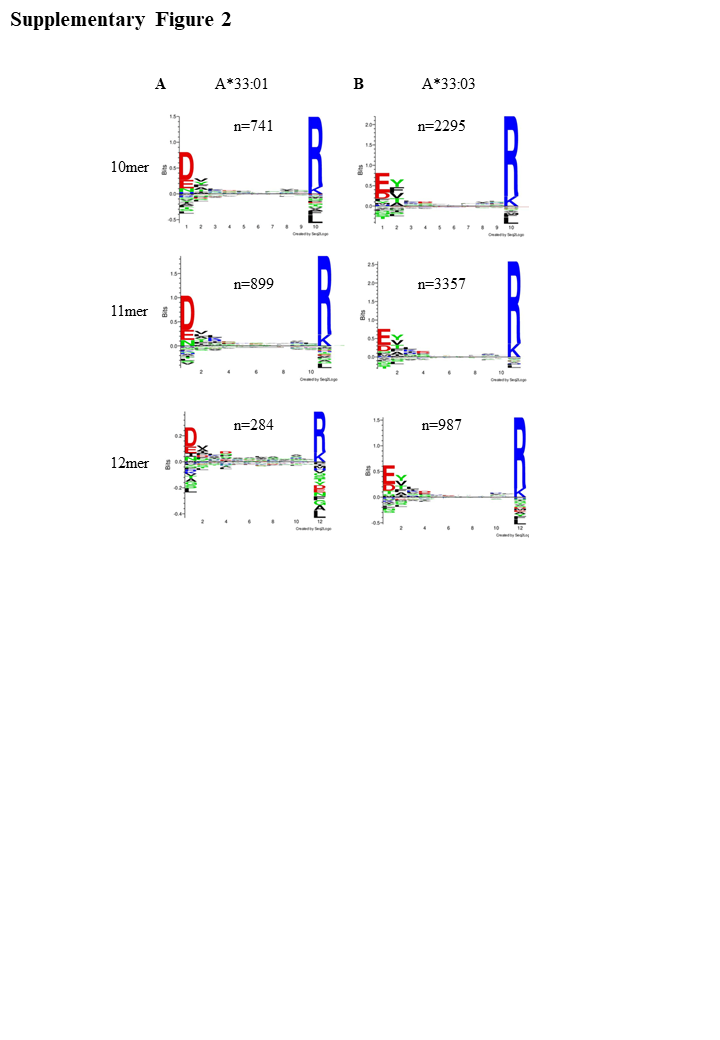

### Figure S3

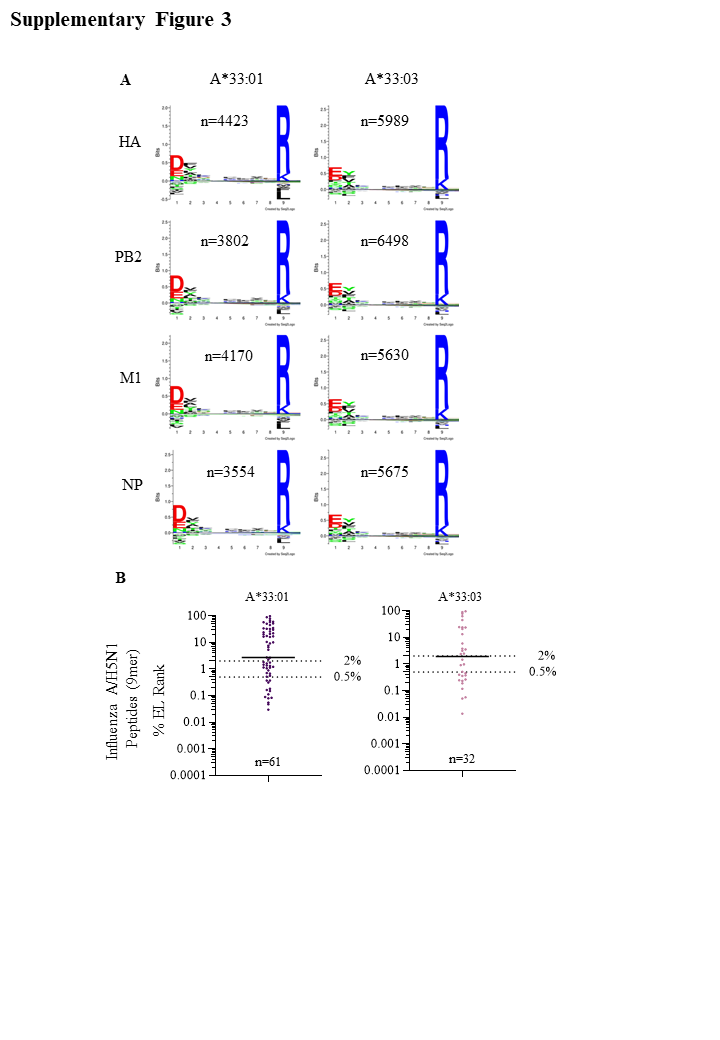

### Figure S4

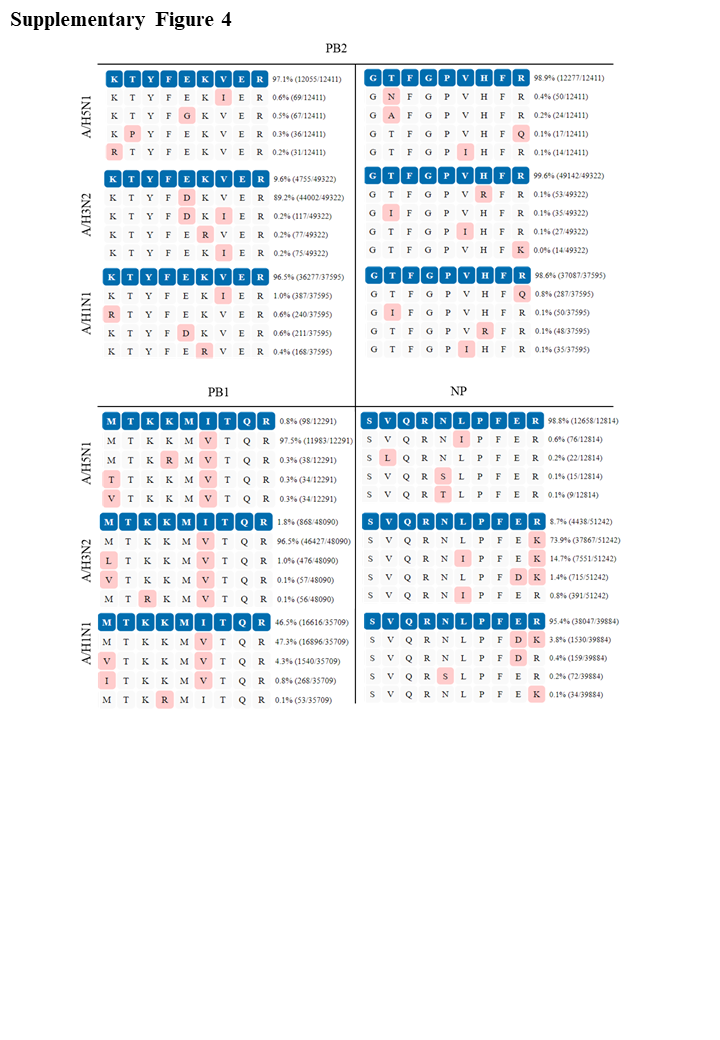
